# Overcoming Daraxonrasib Resistance: Allele-Specific Mechanisms Guide Salvage Therapy in Pancreatic Cancer

**DOI:** 10.64898/2026.07.05.735339

**Authors:** Daniel Dorbin, Juliannie Herrera, Raven Davidson, Naveen Kumar Chandrashekar, Gabrielle Scheuber, Priyanga Jayakrishnan, Christabelle Rajesh, Grayson Johnson, Jason Yuan, Matthew Sochor, John F. Langenheim, Mohammed Aldakkak, Charles Messerly, Jacquelyn Wittmann, Aniko Szabo, Forough Azam Sayahpour, Nathan L. Atallah, Francis C. Peterson, Brian F. Volkman, Mahmoud Ali, Eugene Ke, Douglas B. Evans, Susan Tsai, Nikki K. Lytle, Y. David Seo, Razelle Kurzrock, G. Aaron Hobbs, Mandana Kamgar, Thomas McFall

## Abstract

Clinical-grade RAS inhibitors raise an unresolved question as to whether KRAS-alleles impose constraints on adaptive resistance that can be exploited therapeutically. Using daraxonrasib (RMC-6236), a multi-selective RAS(ON) inhibitor, we compared resistance mechanisms between *KRAS*G12D and *KRAS*G12R, alleles with fundamentally different RAS network dynamics. Daraxonrasib inhibited KRAS^MUT^ primarily through steric occlusion of effector binding, while engaging RAS^WT^ only modestly (∼20%). KRAS^G12R^ is marked by its inability to transactivate RAS^WT^, and it was observed that daraxonrasib resistant KRAS^G12R^ PDAC cells utilize EGFR/RAS^WT^-GTP signaling as the dominant adaptive route. In contrast, KRAS^G12D^ resistance arose through retained KRAS^G12D^-GTP signaling, with a decrease of cyclophilin A (CypA) protein, the binding partner required for daraxonrasib activity. The shift from KRAS^G12R^ dependence to the EGFR/RAS^WT^ conferred sensitivity to trametinib. We confirmed this clinically: a KRAS^G12R^ PDAC patient who progressed after 10 months on daraxonrasib showed intratumoral EGFR/RAS^WT^ activation, and rapid 3D-bioprinted patient-derived toroid modeling predicted sensitivity to trametinib-based combination therapy. Given the aggressive disease trajectory and lack of response to the two immediately preceding lines of therapy, sixth-line trametinib-based combination therapy achieved approximately 5 months of disease control. This patient ultimately achieved 40 months of overall survival, far exceeding the 8–12 month median for metastatic PDAC. Collectively, these data establish a framework in which allele-specific RAS network topology dictates the adaptive resistance landscape, enabling rational selection of targeted therapies with meaningful clinical benefit in metastatic PDAC.

**STATEMENT OF SIGNIFICANCE:** Daraxonrasib resistance mechanisms have allele-specific routes: CypA becomes downregulated in KRAS^G12D^ and reliance on EGFR/RAS^WT^ in KRAS^G12R^. Rapid patient-derived toroids identified sixth-line targeted therapy strategies with an overall survival of 40 months.

## INTRODUCTION

Approximately 90-95% of pancreatic ductal adenocarcinoma (PDAC) cases harbor activating *KRAS* mutations, predominantly affecting codon 12. The most frequent alleles include G12D (35%), G12V (20-30%), and G12R (10-20%), with other rare mutations comprising the remainder (1). Recent therapeutic advances have yielded daraxonrasib (RMC-6236), a multi-selective RAS(ON) inhibitor targeting the active, GTP-bound state of RAS proteins through a tri-complex mechanism involving cyclophilin A (CypA) (2). Unlike allele-specific inhibitors, such as those restricted to *KRAS*G12C (3), daraxonrasib demonstrates broad activity against multiple codon 12 variants (G12D, G12V, G12R, G12A, and G12S), potentially achieving more durable pathway suppression by simultaneously inhibiting both mutant (RAS^MUT^) and wild-type RAS (RAS^WT^) isoforms (4). Phase I clinical trials have established an acceptable safety profile, with reported median progression-free survival (PFS) of 7.6 months in the second-line setting (5). The phase III clinical trial showed significant improvement of outcomes in the second-line metastatic PDAC setting with daraxonrasib as compared to the standard of care chemotherapy, and is expected to lead to eventual FDA approval of this agent (6). However, as with all targeted therapies in metastatic disease, acquired resistance remains inevitable and poses a significant clinical challenge. Treatment strategies following progression on investigational agents are particularly problematic (7); the absence of prospective data governing post-investigational therapy leaves clinicians without evidence-based frameworks to guide management in the late-line setting. While translational models offer valuable insights into resistance mechanisms and therapeutic vulnerabilities, their developmental timeline frequently exceeds the clinical window required for treatment guidance (8).

Resistance to RAS inhibitors is heterogeneous and frequently polyclonal, yet roughly half of patients who progress on KRAS^G12C^ inhibitors lack identifiable co-alterations, leaving the majority without a sequence-based explanation (9,10). This genomic dark fraction likely reflects non-genetic mechanisms; transcriptional reprogramming, lineage plasticity, and adaptive signaling reactivation that are invisible to next-generation sequencing yet functionally decisive. Epithelial-to-mesenchymal transition and histological transformation can drive resistance by inducing a KRAS-independent, mesenchymal cell state (11), underscoring that both protein expression-level and activity-level changes represent a critical and under-characterized axis of therapeutic escape.

The oncogenic impact of mutant *KRAS* is not solely determined by its own constitutive activity, but by how it reshapes the broader RAS network through allele-specific regulation of guanine nucleotide exchange factors (GEFs) and GTPase-activating proteins (GAPs) (12–16). Different *KRAS* mutations impose distinct constraints on GEF and GAP accessibility, thereby differentially modulating the activation state of wild-type RAS paralogs and creating allele-specific network topologies that extend well beyond the mutant allele itself (16,17). Critically, the stoichiometric distribution between mutant and wild-type RAS species is not static; perturbations that shift this balance, such as therapeutic inhibition of KRAS^MUT^, can dynamically redistribute GEF and GAP activity across the network, altering wild-type RAS activation in real time (17). To capture these dynamic network-level effects, we employed an experimental-systems biology framework to interrogate allele-specific RAS network behavior with high temporal resolution, revealing how the stoichiometric interplay between KRAS^MUT^ and RAS^WT^ governs adaptive signaling and shapes the landscape of therapeutic resistance. Together, these systems-level insights informed the construction of allele-specific model systems capable of predicting resistance topology, particularly in cases where resistance emerges in the absence of newly acquired genomic alterations.

Herein, we report a unique study in which a patient with *KRAS*G12R-mutant PDAC achieved an exceptional response to daraxonrasib followed by acquired resistance, providing a rare clinical window into the adaptive biology driven by RAS-targeted therapies. Leveraging preclinical models harboring *KRAS*G12D and *KRAS*G12R alleles, we report a mechanistic basis for *KRAS*G12R hypersensitivity to daraxonrasib and characterize a resistance program defined by a shift toward wild-type RAS (RAS^WT^: H- and NRAS) expression and activity, a G12R-specific adaptive trajectory. Biochemical profiling of the patient’s resected tumor confirmed predominant RAS^WT^ signaling, consistent with our preclinical resistance models, and supported the use of MEK inhibition as part of a rational combination salvage strategy. This hypothesis was supported utilizing multiple model systems, including novel rapid 3D-printed tumor toroids and organotypic *ex vivo*-cultures derived from the patient’s tumor. Off-label trametinib-based combination therapy was associated with radiographic and biomarker evidence of disease control. Stoichiometric analysis of mutant-to-wild-type RAS activity and upstream signaling provided a mechanistic rationale consistent with the observed clinical response, supporting the use of such analyses in the setting of acquired RAS inhibitor resistance. Collectively, these findings reveal that the adaptive landscape of KRAS^G12R^ can confer a therapeutically exploitable vulnerability and establish a framework for mechanism-informed salvage therapy in patients who progress on RAS inhibitors.

## MATERIALS AND METHODS

### Patient and Treatments

Written informed consent was obtained from the patient involved in this manuscript in accordance with the Declaration of Helsinki. The study protocol was reviewed and approved by the institutional review board (PRO00045718 [NCT05630989], PRO00044894 [NCT05802069], and PRO00033119). All procedures were conducted in compliance with recognized ethical guidelines and institutional policies governing human subjects’ research.

### PANC-1 Cell Line Models and Culture

Cell lines were generated by Synthego for the McFall laboratory as described in previous studies. Cells were cultured in DMEM supplemented with 10% FBS, penicillin (100 U/mL), streptomycin (100 μg/mL), and L-glutamine (2 mM) at 37°C with 5% CO_2_. All cell lines were mycoplasma-free (MycoAlert Detection Kit) and used within 11 passages of acquisition to maintain cellular integrity.

### Colorectal Cancer (CRC) Cell Line Library Culture

SW48 cells (RRID:CVCL_LC92) and isogenic counterparts were cultured in Roswell Park Memorial Institute (RPMI) 1640 medium supplemented with FBS (10%), penicillin (100 U/ml), streptomycin (100 μg/ml), and l-glutamine (2 mM). SW403 (RRID:CVCL_0545) cells were cultured in L-15 medium with FBS (10%), penicillin (100 U/ml), streptomycin (100 μg/ml), and l-glutamine (2 mM). LS-180 cells were cultured in Eagle’s minimum essential medium (EMEM) with FBS (10%), penicillin (100 U/ml), streptomycin (100 μg/ml), and l-glutamine (2 mM). LoVo (RRID:CVCL_0399) cells were cultured in F12-K medium with FBS (10%), penicillin (100 U/ml), streptomycin (100 μg/ml), and l-glutamine (2 mM). HCT116 (RRID:CVCL_E4IN) cells were grown in McCoy’s 5A medium with FBS (10%), penicillin (100 U/ml), streptomycin (100 μg/ml), and l-glutamine (2 mM). HCT-15 (RRID:CVCL_0292) cells were grown in RPMI 1640 medium supplemented with FBS (10%), penicillin (100 U/ml), streptomycin (100 μg/ml), and l-glutamine (2 mM). CaCo2 cells were grown in DMEM with FBS (20%), penicillin (100 U/ml), streptomycin (100 μg/ml), and l-glutamine (2 mM). All cells were incubated at 37°C in 5% CO2 (unless cultured in L15 media). SW48 cells were obtained from Horizon Discovery. SW403, HCT116, HCT-15, CaCo2, SW48, and LoVo were obtained from the American Type Culture Collection. Cell lines were *Mycoplasma* free (MycoAlert Mycoplasma Detection Kit) and utilized within 11 passages of acquisition.

### PDAC Cell Line Library Culture

HuP-T3 (RRID:CVCL_1299) cell lines were from the DSMZ German Collection of Microorganisms and Cell Cultures; KP-2 (RRID:CVCL_3004) and TCC-Pan2 (RRID:CVCL_3178) were from the Japanese Collection of Research Bioresources Cell Bank; PK-8 (RRID:CVCL_4718) cell lines were from Riken Cell Bank; the Pa16C cell lines were from Jones et al. (13). HPAC (RRID:CVCL_3517), SW 1990 (RRID:CVCL_1723), PANC-1 (RRID:CVCL_0480), HPAF-II (RRID:CVCL_0313) and PSN1 (RRID:CVCL_1644) cell lines were obtained from the ATCC. HPAC, SW 1990, and PANC-1 cell lines were maintained in DMEM supplemented with 10% FBS without antibiotics. HPAF-II, Pa16c, HuP-T3, KP-2, PK-8, PSN1 and TCC-Pan2 were maintained in RPMI-1640 supplemented with 10% FBS without antibiotics. All cell lines were maintained in a humidified chamber with 5% CO2 at 37°C, *Mycoplasma* free, and used within 11 passages of acquisition.

### Proliferation Assay

Cells (5,000 per well) were seeded in 96-well plates in complete media. Treatments were initiated after the cells were attached (24 hours). At the appropriate time points, cell viability was determined by MTT assay; 5 mg/ml of Dithiothreitol (DTT) MTT in phosphate-buffered saline was added to each well followed by incubation at 37°C for 2 hours. The formazan crystal sediments were dissolved in 100 μl of dimethyl sulfoxide, and absorbance was measured at 590 nm using a Tecan Infinite 200 PRO plate reader (RRID:SCR_020543). Each treatment was performed in eight replicate wells and repeated three different times.

### Colony Formation Assay

Cells were trypsinized, and 1000 cells per well were plated in triplicate six-well (60 mm) plates in either complete media (10% FBS) or serum reduced media (2% FBS). Colonies were formed after at least 4 weeks. Cells were fixed with ice-cold methanol and stained with crystal violet. Images were obtained using the Li-Cor Odyssey M imaging station (RRID:SCR_025709) under the 700 nM reading. Three experimental replicates were performed.

### siRNA Knockdown

Cells were plated in adherent culture plates containing siRNA and Lipofectamine RNAiMAX (13778150, Thermo Fisher) at 10 pmol siRNA per 10,000 cells in Opti-MEM reduced medium. siRNAs used were KRAS siRNA (S120703), HRAS siRNA (S807, Thermo Fisher), NRAS siRNA (S55, Thermo Fisher) and control nontargeting siRNA (AM4611, Thermo Fisher).

### Western Blot Analysis

Cell lysates were generated using lysis buffer (1862301, Thermo Fisher Scientific) containing a protease inhibitor cocktail (Cell Signaling Technology) and incubated on ice for 1 hour with brief vortexing every 5 minutes. Total protein concentration was determined using the Pierce protein assay (Thermo Fisher Scientific). Protein samples were resolved on 12% SDS-PAGE or 4%-18% gradient gels and transferred to PVDF membranes (Millipore) for 30 minutes at 25 V using the Trans-Blot Turbo Transfer System (Bio-Rad, RRID:SCR_023156). Blots were probed with the indicated primary and fluorophore-conjugated secondary antibodies and visualized using the Li-Cor Odyssey M imaging station (RRID:SCR_025709). Comparative changes were measured with Li-Cor Image Studio software from three independent experiments (n = 3), normalized to the indicated loading control. Primary antibodies: mouse anti-pERK (BioLegend 675502, RRID:AB_2565604), rat anti-ERK (BioLegend 686902, RRID:AB_2629535), mouse anti-MEK (CST 9122, RRID:AB_823567), rabbit anti-pMEK (CST 9154, RRID:AB_2138017), rabbit anti-pS473-AKT (CST 4060), rabbit anti-pT308-AKT (CST 9275, RRID:AB_329828), mouse anti-AKT (CST 58295, RRID:AB_2799545), anti-KRAS (Sigma-Aldrich WH0003845M1, RRID:AB_1842235), mouse anti-NRAS (Abcam ab188369, RRID:AB_2934137), rabbit anti-HRAS (Abcam ab32417, RRID:AB_726959), and mouse anti-GAPDH (Santa Cruz sc-47724).

### EGFR ELISA

EGFR expression was quantified in 25,000 cells per line using a commercial human soluble EGFR ELISA kit (ab269558) per manufacturer instructions, with absorbance read at 450 nm on a Tecan SPARK reader (RRID:SCR_021897).

### Phosphorylated EGFR ELISA

EGFR phosphorylation at tyrosine 1068 was quantified using a cell-based ELISA (Human EGFR [Phospho] [pY1068] Kit, Thermo Fisher KHR9081) per manufacturer instructions, with 25,000 cells per line and absorbance read at 450 nm (Tecan SPARK, RRID:SCR_021897).

### RAS-GTP Assay (SDS-PAGE Analysis)

GTP-bound RAS isoforms were captured with the Active Ras Pull-Down and Detection Kit (Thermo Fisher) per manufacturer instructions. RBD-enriched lysates were probed with mouse anti-KRAS (Sigma-Aldrich WH0003845M1), rabbit anti-NRAS (Proteintech CL650-10724), rabbit anti-HRAS (Abcam ab32417), and mouse anti-GAPDH (Santa Cruz sc-47724). Total lysates were probed with anti-pERK (BioLegend 675502) and anti-total ERK (BioLegend 686902).

### Two-Dimensional Gel Electrophoresis

Lysates were processed using the ReadyPrep 2-D Cleanup Kit (Bio-Rad). For isoelectric focusing, 50 μg protein samples were resuspended in rehydration solution (8 M urea, 2 M thiourea, 4% CHAPS, 0.5% IPG buffer pH 3-10, 1 mM DTT). First-dimension separation used Criterion IEF gels (pH 3-10) with a stepped voltage protocol (100 V/60 min, then 250 V/60 min). Strips were equilibrated, then resolved on 12% polyacrylamide Criterion gels (120 V, ∼2 h) and transferred to PVDF membranes. Membranes were probed with pan-RAS antibody (Thermo Fisher Active Ras kit) and secondary antibody (Novus NBP1-72928) and imaged on a LI-COR Odyssey M (RRID:SCR_025709).

### One-Dimensional Isoelectric Focusing of GTP-Loaded and Total RAS Isoforms

Cells were harvested by scraping into extraction buffer (25 mM Tris-HCl pH 7.2, 150 mM NaCl, 5 mM MgCl_2_, 1% NP-40, 5% glycerol) and processed for RBD affinity precipitation. RBD-bound proteins were resolved on 12% SDS-PAGE; 21-kDa bands were excised and recovered using the ReadyPrep 2-D Cleanup Kit. Purified samples were focused on Criterion Bio-Lyte IEF gels (pH 3-10) using a two-step voltage gradient (100 V/60 min, 250 V/60 min), transferred to PVDF, and probed with pan-RAS antibody and anti-mouse DyLight 800 secondary (Novus NBP1-72928). Quantification used LI-COR Image Studio (RRID:SCR_015795), normalized to input controls. Proportion of occlusion was calculated as signal(RBD-PD) − signal(RBD+Cyclosporine); the amount of GTP hydrolysis equaled signal(RBD+Cyclosporine).

### CypA RAS-GTP Assay (SDS-PAGE Analysis)

GTP-bound RAS isoforms were captured using GST-tagged CypA. PANC-1 isogenic cells were lysed in lysis/binding/wash buffer (25 mM Tris-HCl pH 7.2, 150 mM NaCl, 5 mM MgCl_2_, 1% NP-40, 5% glycerol) with protease inhibitors, clarified at 16,000 × g, and quantified by Pierce 660 assay. GST-CypA fusion protein (80 μg) with 1 μM RMC-6236 or DMSO was bound to glutathione resin, washed, and eluted in 2× Laemmli buffer.

### Tumor Xenografts

Studies were approved by the MCW Animal Resources Department and IACUC (protocol AUA00008123). NOD-SCID IL2Rγ−/− mice (RRID:IMSR_NM-NSG-021, Jackson Laboratories #005557) received subcutaneous injections of 1 × 10^6^ cells into the right flank. Tumor volumes were calculated as V = ½ (Length × Width^2^). Cohorts included both sexes distributed equally among arms; pooled analyses showed no sex-based differences. Animals were housed under specific-pathogen-free conditions (12-hour light/dark, 23°C, 30-70% humidity) with standard chow and water ad libitum.

### Toroid Printing

Adherent cells (1.5 × 10^6^) were seeded and, upon reaching 70% confluency, magnetized with NanoShuttle-PL (Greiner Bio-One 657841) for up to 16 hours, then dissociated with TrypLE Express (Gibco). For patient tumors, dissociated cells were pelleted, resuspended with NanoShuttle-PL, and processed through three cycles of centrifugation/resuspension. Magnetized cells were seeded in cell-repellent plates with a levitation drive (Greiner Bio-One), aggregated, then transferred to 96-well cell-repellent microplates with a ring drive for toroid formation. Toroids were imaged at 20 μm resolution (Odyssey M, LI-COR). Compounds were added at 10× final concentration, with imaging at 24 and 48 hours post-treatment.

### Toroid Quantification

Microscopy images were analyzed with custom Python scripts (v3.12.7); wells were cropped using the Python Imaging Library (v10.4.0). Feature representations were generated with a ResNet-50 CNN pretrained on ImageNet (18), implemented via HuggingFace transformers (v4.48.2) on PyTorch (v2.6.0+cu126, NVIDIA RTX 4050, CUDA 12.7.33). Four rotational variants (0°, 90°, 180°, 270°) were averaged per well for rotational invariance. Degradation was quantified as the cosine similarity (NumPy v1.26.4) between each timepoint’s encoding vector and the baseline vector, with replicate scores normalized to a 0-1 range. Code is available on GitHub (see Code Availability).

### Ex Vivo Organotypic Culture Assays

Image-guided core biopsies (MCW Surgical Oncology Tissue Bank, IRB PRO00033119) were transported in cold RPMI 1640 with 10% FBS/1% Pen-Strep. Within 15-30 minutes, 3-4 mm slices were placed on 0.4 mm membrane inserts (Millicell-CM) in RPMI 1640/10% FBS/1% Pen-Strep, cultured at 37°C/5% CO2 with media changes every 2-3 days. After 24 hours, slices were treated for 24-48 hours before MTS viability assay and fixation for H&E/IHC. Slice area was quantified in ImageJ. FFPE sections were H&E stained; nuclear density, tumor content, and tumor-stroma ratios were quantified with QuPath. IHC primary antibodies: cleaved Caspase-3 (R&D MAB835-SP), Ki-67 (Santa Cruz sc-23900), CD45 (Santa Cruz sc-1187), EpCAM (Santa Cruz sc-25308), with Vector ImmPress HRP secondary and DAB development. Positive cells were quantified in QuPath by three independent observers with consensus resolution.

### DNA Isolation and Whole Exome Sequencing

Genomic DNA was isolated from PANC-1 lines (G12D, G12R, G12D-RR, G12R-RR) using the GeneJET Genomic DNA Purification Kit (Thermo Fisher). Exome sequencing was performed by Novogene using the Agilent SureSelect Human All Exon V6 kit, sequenced on a NovaSeq X Plus (150 bp paired-end).

### Patient Genomic Profiling

Somatic alterations were evaluated by the CLIA-certified, CAP-accredited Tempus xT assay (Tempus AI, Inc.), a 648-gene next-generation sequencing panel. Variants identified at diagnosis and at progression on daraxonrasib were compared to assess for acquired co-alterations.

### CypA Restoration in Daraxonrasib-Resistant Cell Lines

G12D-RR cells were transfected with pcDNA encoding full-length human CypA using Lipofectamine 2000; restoration was confirmed by western blot. An identical approach was applied to G12R-RR cells as a control. Cells were maintained in daraxonrasib-containing media and assessed for KRAS-GTP, pERK, and proliferation.

### Immunohistochemical Analysis of Patient Tumor EGFR Expression

EGFR expression in the resected metastatic tumor was assessed by IHC at MCW Pathology Services using an anti-EGFR antibody (LSBio LS-C95382-1, RRID:AB_1931104) per standard clinical protocol; positivity was scored by percentage membranous staining and intensity by a board-certified pathologist.

### CellTiter-Glo Luminescent Viability Assay

Toroids were gently disrupted and transferred to opaque 96-well plates. CellTiter-Glo 3D reagent (Promega G9681) was added 1:1 with media, incubated 10 minutes, and luminescence measured (Tecan SPARK, RRID:SCR_020543), normalized to vehicle-treated wells.

### Live-Dead Cell Analysis

Dissociated toroid cells were mixed 1:1 with 0.4% trypan blue and quantified with a Bio-Rad TC20 Automated Cell Counter (RRID:SCR_025462) in triplicate.

### Quantification and Statistical Analysis

The number of biological replicates is denoted by “n” and the number of experimental replicates is specified in each figure legend. Data are reported as mean ± standard deviation and analyzed using GraphPad Prism 8 (RRID:SCR_000306). Two-condition comparisons used an unpaired two-tailed t-test (equal variances assumed) from at least three independent experiments; comparisons of three or more groups used one-way ANOVA with post hoc Tukey’s test. Significance is reported as P < 0.05, 0.01, 0.001, 0.0001 or exact values, indicated in each figure, with the statistical test noted in the corresponding legend.

## RESULTS

### KRAS^G12R^ Exhibits Attenuated RAS Network Signaling and Enhanced Sensitivity to RAS Inhibition

KRAS^G12R^ displays distinct biochemical signaling compared to KRAS^G12D^, with reduced overall RAS pathway output (13). We therefore used PANC-1 isogenic cell lines harboring *KRAS*G12R or *KRAS*G12D (13). KRAS^G12R^ cells showed markedly reduced MEK and ERK phosphorylation across three independent clones **(Fig. 1A-B)**. Although KRAS^G12R^ shows reduced GTP hydrolysis and exchange rates relative to KRAS^G12D^ (19), it cannot transactivate RAS^WT^ isoforms (HRAS and NRAS), yielding paradoxically lower total RAS pathway activation despite elevated KRAS^MUT^ activity (13). RBD-IEF assays (14–16) confirmed reduced HRAS^WT^ and NRAS^WT^ activation in *KRAS*G12R cells **(Fig. 1C-D)**. Consistent with this, *KRAS*G12R xenografts showed delayed growth kinetics **(Fig. 1E and S1A)**, aligning with the improved clinical outcomes and lower ERK signaling potential reported for KRAS^G12R^ PDAC (13,20,21).

**Figure 1.**
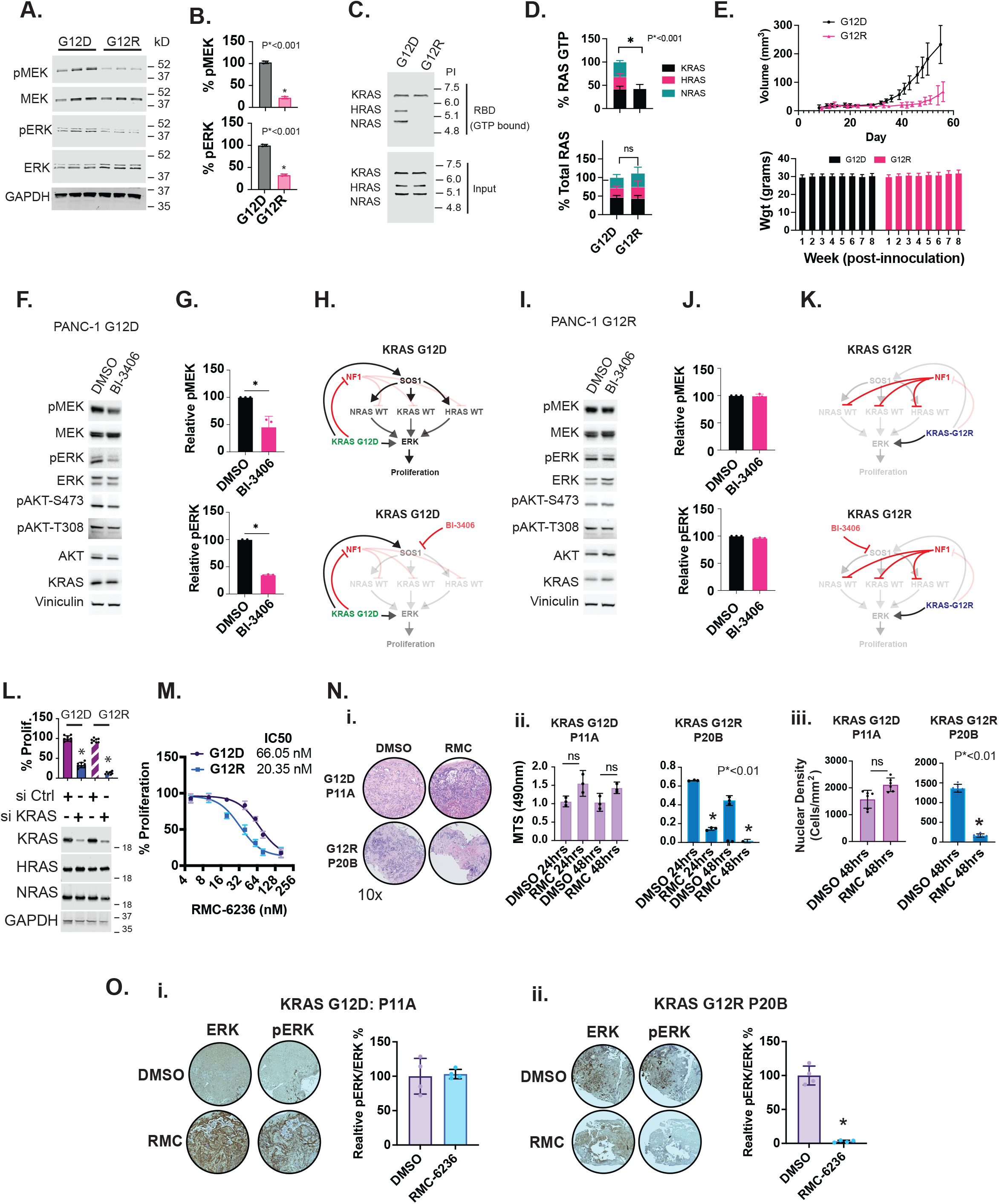
Molecular Signaling Profile of KRAS ^G12R^ and KRAS^G12D^ and Therapeutic Vulnerability to KRAS^MUT^ Inhibitors. **A**. PANC-1 isogenic cells harboring KRAS^G12D^ or KRAS^G12R^ were analyzed by SDS-PAGE and immunoblotted for phosphorylated MEK (pMEK), total MEK, phosphorylated ERK (pERK), and total ERK, with GAPDH as a loading control. **B**. Quantification across three independent experiments demonstrated reduced pMEK and pERK in PANC-1 G12R cells. **C**. Total RAS protein (input) and active RAS-GTP (isolated by RBD pull-down assay using the RAF RAS-binding domain) were separated by isoelectric focusing and detected with a pan-RAS antibody. **D**. Fluorescence quantification of both input lysates (bottom) and RBD-captured RAS-GTP (top) revealed reduced activation of HRAS and NRAS in PANC-1 KRAS G12R cells compared to KRAS G12D cells. **E**. Xenografts of PANC-1 isogenic cell lines in NOD-SCID IL2Rg^-/-^ (NSG) mice (n=5) with growth curves over 58 days. Data points represent mean values, and error bars indicate standard deviation. Graph is representative of three separate experiments (Top). Mouse weight was measured over the course of the study, and no difference was observed over the course of the study (Bottom). **F**. PANC-1 KRAS^G12D^ cells (in complete media) were treated with vehicle (DMSO) or 10 µM BI-3406 for 12 hours. Cells were lysed, and proteins were resolved by SDS-PAGE and immunoblotted for RAS pathway proteins and their phosphorylated (activated) forms. *G*. Quantification of pMEK and pERK from three independent experiments shown in panel F. **H**. Proposed molecular mechanism of RAS^WT^ activation in the KRAS^G12D^ context. **I**. PANC-1 KRAS^G12R^ cells (in complete media) were treated with vehicle (DMSO) or 10 µM BI-3406 for 12 hours. Cells were lysed, and proteins were resolved by SDS-PAGE and immunoblotted for RAS pathway proteins and their phosphorylated (activated) forms. **J**. Quantification of pMEK and pERK from three independent experiments shown in panel I. **K**. Proposed molecular mechanism of why altering SOS activity does not alter ME/ERK signaling in the KRAS^G12R^ context. **L**. PANC-1 isogenic cell lines were treated with non-targeting control siRNA or KRAS-targeting siRNA. At 24 hours post-transfection, whole-cell lysates were subjected to SDS-PAGE and immunoblotted with antibodies specific for KRAS, HRAS, and NRAS (bottom panel). Cell viability was assessed in parallel using MTT assay at 72 hours post-transfection (top panel). **M**. PANC-1 isogenic cell lines were treated with increasing concentrations of RMC-6236, and cell viability was assessed after 5 days using MTT assay to determine IC_50_. **N**. Organotype Ex-vivo slice cultures from treatment-naïve KRAS^G12D^ (Patient P11A) and KRAS^G12R^ (Patient P20B) PDAC patient biopsies were treated with DMSO (vehicle control) or 200 nM daraxonrasib to assess acute drug sensitivity. (i) Hematoxylin and eosin (H&E) staining confirming tumor tissue. (ii) MTS viability assay showing relative proliferation at 24 and 48 hours post-treatment. (iii) Quantification of nuclear density changes at 48 hours. **O**. Following analysis of cell density and growth shown in **Fig. 1N**, IHC for pERK and ERK was performed on tissue sections, and the relative pERK/ERK signal ratio was quantified across four independent tissue sections for Patient P11A (i) and Patient P20B (ii). All experiments were performed at least three times for statistical rigor. Statistical differences were calculated by two-tailed unpaired t-test and P values are indicated (for two groups). For experiments with three or more groups statistical comparisons were calculated by one-way ANOVA followed by post hoc Tukey’s test for multiple comparisons, and P values are indicated in the panels. Each experiment is representative of three experiments; quantifications represent mean and error bars represent standard deviations from three separate experiments

RAS^WT^ transactivation in KRAS^G12D^ PDAC has been attributed to two mechanisms: KRAS^G12D^ acting as a stoichiometric sink for neurofibromin (NF1), a GAP that normally inactivates RAS^WT^, and allosteric activation of SOS1, a GEF that promotes RAS^WT^-GTP loading (13). Both mechanisms are absent in the context of *KRAS*G12R (13,14,22). To test this orthogonally, we treated *KRAS*G12D cells with the SOS1 inhibitor BI-3406 and observed decreased pMEK and pERK **(Fig. 1F-H)**, indicating SOS1-dependent ERK signaling; no such effect occurred in KRAS^G12R^ cells **(Fig. 1I-K)**, consistent with the absence of SOS1 feedforward activation in this context. As PANC-1 cells express minimal RTKs (13), we confirmed these SOSi effects were RTK-independent by repeating the experiment in serum-free media with identical results **(Fig. S1B-C)**. SOS1 inhibition similarly decreased pMEK/pERK in *KRAS*G12D lines (ASPC1, SW1990; **Fig. S1D**) but not in *KRAS*G12R lines (PK8, PSN1, KP2; **Fig. S1E**), recapitulating the isogenic findings.

The network hypothesis predicts that SOS1 inhibition reduces RAS^WT^-GTP (HRAS, NRAS) in *KRAS*G12D cells by disrupting SOS1-mediated transactivation, an interaction sterically blocked by the G12R substitution (13,22). In serum-deprived SW1990 (*KRAS*G12D) cells, basal HRAS-GTP and NRAS-GTP levels were elevated relative to TccPan-2 (KRAS^G12R^) cells and decreased with BI-3406 treatment, whereas TccPan-2 HRAS-GTP/NRAS-GTP levels were lower at baseline and unaffected by BI-3406 **(Fig. S1F)**. Combined, these data support a KRAS allele-specific mechanism where G12D, but not G12R, promotes activation of RAS^WT^ alleles through a SOS1-dependent interaction.

This reliance on KRAS^MUT^ signaling suggests hypersensitivity to KRAS-targeted inhibition in the *G12R* context. Indeed, genetic KRAS knockdown suppressed proliferation more effectively in G12R-expressing cells than G12D cells **(Fig. 1L)**, and *KRAS*G12R cells showed a lower daraxonrasib IC50 **(Fig. 1M)**. To validate this in a translational model, we performed *ex vivo* organotypic culture assays **(Fig. S2A)** on treatment-naive biopsies from two patients: P11A (*KRAS*G12D) and P20B (*KRAS*G12R) **(Fig. 1N: i)**. Daraxonrasib treatment produced significantly greater viability reduction and nuclear density loss in the *KRAS*G12R slices than the *KRAS*G12D slices **(Fig. 1N: ii-iii)**. IHC corroborated this divergence: P11A (*KRAS*G12D) showed modest cleaved caspase induction and robust CD45 upregulation with unchanged Ki-67/EpCAM **(Fig. S3B-C)**, whereas P20B (*KRAS*G12R) showed robust cleaved caspase staining with minimal CD45 change and no Ki-67/EpCAM alteration **(Fig. S3D-E)**, consistent with the enhanced sensitivity in **Fig. 1L-M**. pERK/ERK IHC on the same slices after 48 hours showed no change in pERK for P11A **(Fig. 1O-i)**, suggestive of a temporally delayed response in the G12D setting, but marked reduction in P20B **(Fig. 1O-ii)**, consistent with effective pathway suppression and corroborating the proliferative data in **Fig. 1L-M**.

### Daraxonrasib Exhibits Isoform-Selective Inhibition, Preferentially Targeting KRAS^MUT^ over RAS^WT^

Daraxonrasib inhibits RAS by recruiting CypA to GTP-bound RAS, forming a tri-complex that sterically occludes effector binding and accelerates GTP hydrolysis (23). Chemical-biology studies suggested preferential engagement with KRAS^MUT^ over RAS^WT^ (23), but in-cell selectivity was unclear. The intratumoral concentration to inhibit growth is predicted to be 80 nmol/L (2), which effectively suppressed proliferation in our growth assays **(Fig. 1M)**. To quantify selectivity, we performed RAS-GTP pulldown assays (RBD-GST) in PANC-1 isogenic lines followed by IEF to resolve individual RAS isoforms at 80 nM (14–16). Daraxonrasib effectively blocked KRAS^MUT^-GTP binding to RBD, confirming robust mutant KRAS inhibition, while RAS^WT^ (HRAS, NRAS) RBD binding remained largely intact in *KRAS*G12D cells, indicating continued RAS^WT^ effector engagement **(Fig. 2A: i and 2C: i)**. RBD binding quantification showed near-complete KRAS^G12D^ inhibition (∼90-95%) but minimal RAS^WT^ inhibition (∼10-20%) **(Fig. 2B: i)**. In *KRAS*G12R cells, which cannot directly transactivate RAS^WT^ via SOS1 activation, RAS^WT^-GTP was virtually undetectable before or after daraxonrasib treatment, serving as an internal negative control **(Fig. 2A: i and 2B: ii)**.

**Figure 2.**
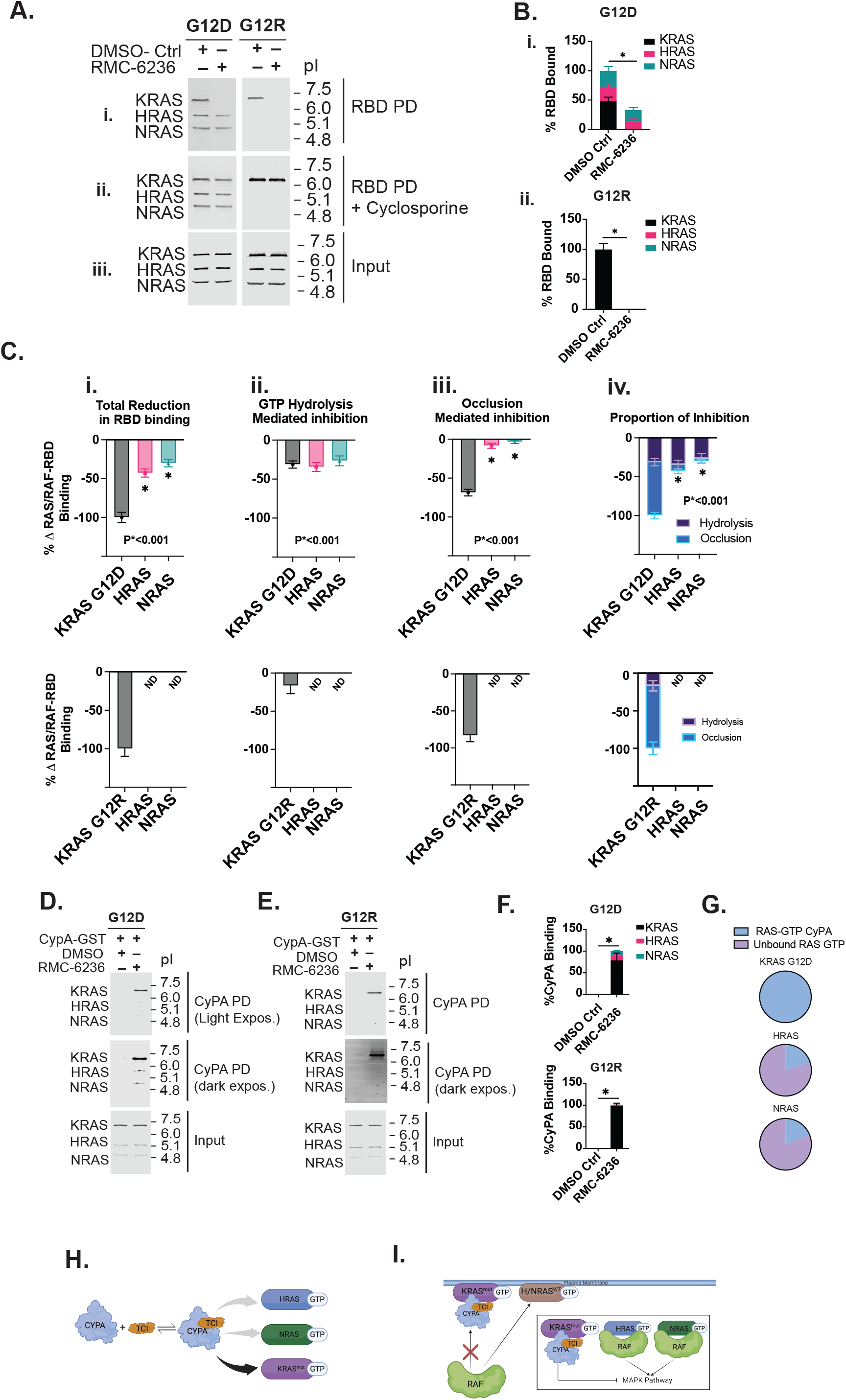
Daraxonrasib Exhibits Preferential Binding to KRAS^MUT^ Over RAS^WT^. **A**. PANC-1 G12D and G12R cells were treated with 80 nM RMC-6236 for 24 hours and subjected to two distinct immunoprecipitation approaches to assess RAS-GTP binding: (i) RBD pull-down was performed using standard lysis and wash buffers that preserve cyclophilin A (CyPA) interaction with RAS-GTP. Captured proteins were separated by isoelectric focusing, transferred to PVDF membrane, and probed with pan-RAS antibody. (ii) RBD pull-down was performed in the presence of cyclosporine A to disrupt CyPA binding and enable direct RBD interaction with RAS-GTP. (iii) Whole cell lysates for input were resolved by isoelectric focusing (IEF). **B**. Quantification from three separate experiments from panel A:(i) of RAS-GTP isoforms binding to RBD in Vehicle or RMC-6236 treated cells. **C**. Quantitative analysis derived from three independent experiments shown in panel A, illustrating: (i) total reduction in RAS isoform binding to RBD, (ii) signal loss attributable to GTP hydrolysis, (iii) signal loss due to CyPA-mediated steric occlusion of the RBD-binding interface, and (iv) combined proportion of RAS-GTP signal loss resulting from both CyPA occlusion and GTP hydrolysis. **D**. PANC-1 G12D cells were subjected to CypA-GST pulldown with equimolar RMC-6236 or DMSO control. Captured RAS-GTP was resolved by one-dimensional IEF. **E**. PANC-1 G12R cells were subjected to CypA-GST pulldown with equimolar RMC-6236 or DMSO control. Captured RAS-GTP was resolved by one-dimensional IEF. **F**. Quantification of each RAS isoform binding to CypA was calculated from three independent experiments. **G**. Proportion of each RAS/CypA bound/unbound for PANC-1 KRAS G12D cells. **H**. CyPA/RMC-6236 complex has a selective bias toward binding KRAS^MUT^-GTP over RAS^WT^-GTP. **I**. Proposed resistance mechanism: RAS^WT^ isoforms bypass daraxonrasib/CypA complex inhibition due to reduced binding affinity. All experiments were performed three times. Statistical differences were calculated by two-tailed unpaired t-test and P values are indicated (for two groups). For experiments with three or more groups statistical comparisons were calculated by one-way ANOVA followed by post hoc Tukey’s test for multiple comparisons, and P values are indicated in the panels. Each experiment is representative of three experiments; quantifications represent mean and error bars represent standard deviations from three separate experiments.

Parallel cyclosporine-competition experiments, which disrupt the daraxonrasib/CypA/RAS tri-complex **(Fig. 2A: ii)**, alongside standard RBD assays **(Fig. 2A: i)**, showed that KRAS^MUT^ inhibition occurs primarily through steric occlusion (∼75-80% of total inhibition; **Fig. 2C: i, iii, iv**), with accelerated GTP hydrolysis contributing the remainder (∼20-25%; **Fig. 2C: ii**). The limited RAS^WT^ inhibition occurred predominantly through enhanced hydrolysis rather than steric blockade **(Fig. 2C: i-iv)**.

CypA-GST pulldown assays with equimolar daraxonrasib confirmed direct tri-complex formation: the complex bound KRAS^MUT^-GTP strongly (normalized to 100%) but less than 20% of RAS^WT^-GTP in *KRAS*G12D cells **(Fig. 2D and 2F: Top)**. In *KRAS*G12R cells, which lack basal RAS^WT^ activation **(Fig. 1C and 1D)**, RAS^WT^-GTP remained virtually undetectable regardless of treatment **(Fig. 2E and 2F: Bottom)**. Notably, when robust RAS^WT^-GTP was present, CypA/daraxonrasib binding was under 25% **(Fig. 2G)**.

This selectivity for KRAS^MUT^ may explain the differential drug sensitivity between *KRAS*G12R and *KRAS*G12D cells **(Fig. 1M)**: KRAS^G12R^ cells rely predominantly on KRAS^MUT^ signaling due to their inability to transactivate RAS^WT^ **(Fig. 1C)** (13), conferring enhanced daraxonrasib sensitivity, whereas the gain-of-function properties of KRAS^G12D^ drive robust RAS^WT^ transactivation that is refractory to inhibition, necessitating higher drug concentrations to achieve equivalent pathway suppression **(Fig. S3A)**. This persistent RAS^WT^-GTP likely reflects the drug-free fraction of mutant KRAS retaining scaffolding capacity that sustains SOS1-mediated exchange and NF1 sequestration (13–16,22). Because the CypA-daraxonrasib complex does not engage RAS^WT^ equivalently, preserved RAS^WT^-driven signaling could have direct implications for the development of adaptive resistance in tumors **(Fig. 2H-I)**.

### Temporal Dynamics of Adaptive Resistance Reveal Specific Stoichiometric Remodeling

To model adaptive resistance under physiologically relevant conditions, we cultured PANC-1 G12R and G12D isogenic lines at their IC50 for 14 days, with drug replenished every three days, and analyzed lysates by western blot for RAS isoforms, EGFR, total ERK, and pERK. PANC-1 KRAS^G12D^ cells showed transient upregulation of RAS^WT^ isoforms by day 4 that declined below baseline by day 14, while KRAS^G12D^ protein and pERK remained elevated throughout, reaching 1.5-fold above baseline by day 14 **(Fig. 3A and B)**, indicating an initial RAS^WT^ compensation phase followed by stable resistance through sustained KRAS^MUT^ expression. PANC-1 KRAS^G12R^ cells instead showed a switch from KRAS^MUT^ to an EGFR/RAS^WT^ expression program: all three RAS isoforms increased by day 2 with pERK still suppressed, EGFR became detectable by day 3 and rose through day 10 while KRAS^G12R^ declined below baseline by day 14 and HRAS/NRAS stabilized approximately 2-fold above baseline, with pERK equilibrating to roughly 50% of baseline **(Fig. 3C and D)**.

**Figure 3.**
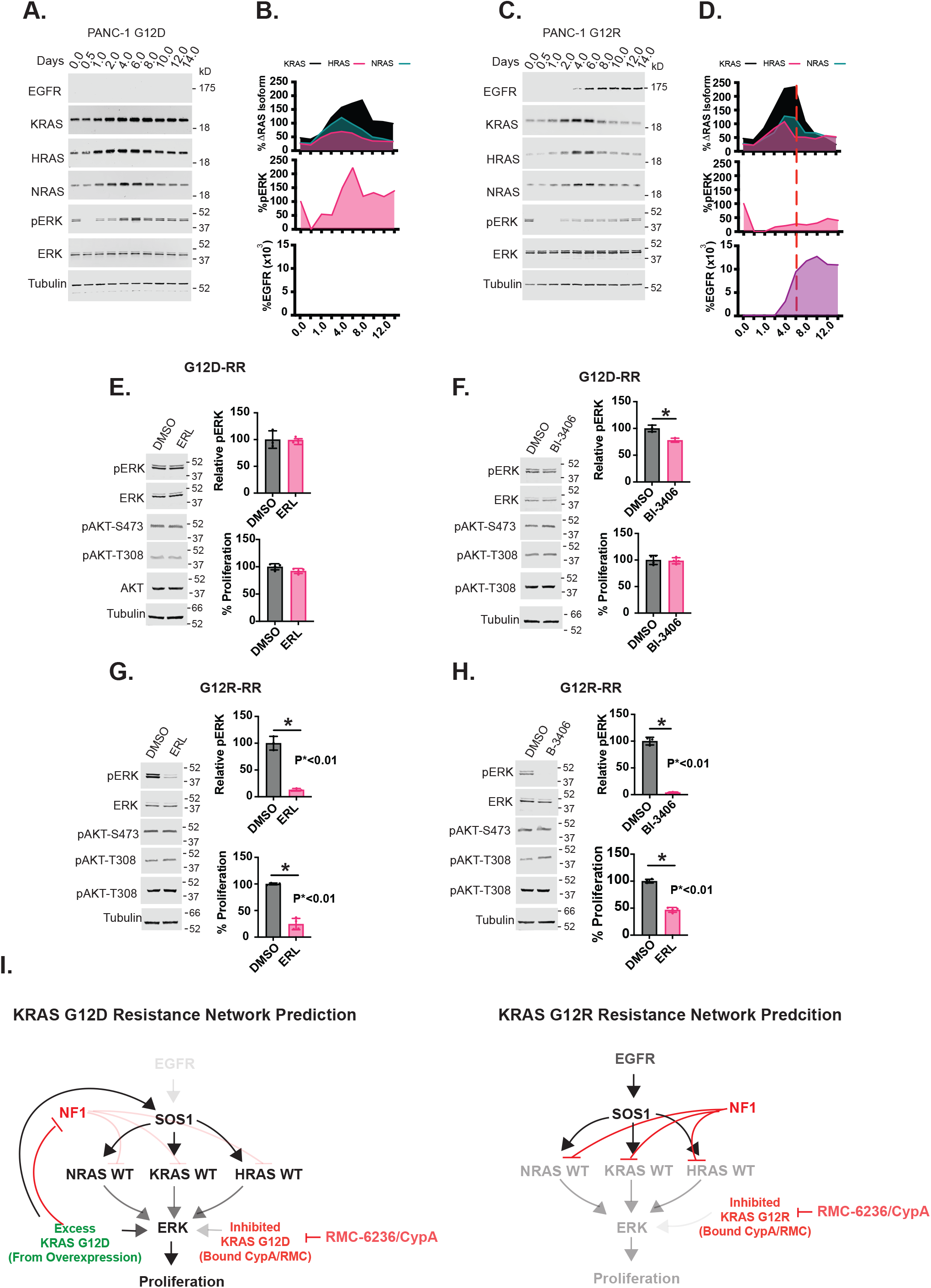
Mutant KRAS inhibition results in temporal compensation by alternative RAS isoforms. **A**. PANC-1 G12D cells were treated with vehicle control (0 nM) or 60 nM RMC-6236 for the indicated times. Cell lysates were analyzed by western blot for EGFR, KRAS, HRAS, NRAS, phosphorylated ERK (pERK), total ERK, and tubulin (loading control). **B**. quantification of fluorescent signal for proteins from **panel A**: RAS isoform expression relative to baseline, normalized to the baseline stoichiometry in PANC-1 G12D cells (top). Quantification of pERK/total ERK ratio over time (middle). Quantification of EGFR expression levels over time (bottom). **C**. PANC-1 G12R cells were treated with vehicle control (0 nM) or 20 nM RMC-6236 for the indicated times. Cell lysates were analyzed by western blot for EGFR, KRAS, HRAS, NRAS, pERK, total ERK, and tubulin (loading control). **D**. Quantification of fluorescent signal for proteins from **Panel C**: RAS isoform expression relative to baseline, normalized to the baseline stoichiometry in PANC-1 G12R cells (top). Quantification of pERK/total ERK ratio over time (middle). Quantification of EGFR expression levels over time (bottom). **E**. PANC-1 G12D-RR cells maintained in RMC-6236–containing media were treated with erlotinib (200 nM, 24 hours). Cell lysates were resolved by SDS-PAGE and immunoblotted for pERK, total ERK, phosphorylated AKT (pAKT) and total AKT. (Right) Quantification from three separate experiments showing changes in pERK levels (Left). **F**. PANC-1 G12D-RR cells maintained in RMC-6236–containing media were treated with BI-3406 (10 uM, 24 hours). Cell lysates were resolved by SDS-PAGE and immunoblotted for pERK, total ERK, pAKT and total AKT. Quantification from three separate experiments showing changes in pERK levels (Right). **G**. PANC-1 G12R-RR cells maintained in RMC-6236–containing media were treated with erlotinib (200 nM, 24 hours). Cell lysates were resolved by SDS-PAGE and immunoblotted for pERK, total ERK, pAKT and total AKT. Quantification from three separate experiments showing changes in pERK levels. **H**. PANC-1 G12R-RR cells maintained in RMC-6236–containing media were treated with BI-3406 (10 uM, 24 hours). Cell lysates were resolved by SDS-PAGE and immunoblotted for pERK, total ERK, pAKT and total AKT. Quantification from three separate experiments showing changes in pERK levels. **I**. Network systems hypothesis for ERK reactivation under RMC-6236 treatment, incorporating allosteric SOS1 activation, NF1 inhibition, EGFR-mediated SOS1 activation, and wild-type RAS escape from RMC-6236 engagement.

To establish stable resistant models, we exposed isogenic lines to continuous IC50 daraxonrasib for several months without dose escalation, then treated the resulting -RR derivatives with erlotinib (EGFR inhibitor) or BI-3406 (SOS1 inhibitor). KRAS^G12D^-RR cells showed no change in pERK or proliferation with erlotinib, consistent with their lack of EGFR expression **(Fig. 3E)**, and only a modest pERK decrease with BI-3406, with no effect on proliferation **(Fig. 3F)**, possibly reflecting the intrinsic nucleotide exchange rate of KRAS^G12D^ (24) and residual SOS1 activity by free KRAS^G12D^-GTP (13,22); consistent with PI3K not being a dominant RAS effector in PDAC (21), pAKT was unaffected by either drug.

In contrast, KRAS^G12R^-RR cells showed robust pERK reduction and significantly decreased proliferation with either erlotinib or BI-3406 **(Fig. 3G and 3H)**, indicating EGFR and RAS^WT^ dependence. Applying our systems network hypothesis (13,14,16,17,25), we propose that G12D-RR cells sustain resistance via retained KRAS^G12D^-GTP that drives ERK signaling, with residual RAS^WT^ activity further augmented by allosteric SOS1 activation and NF1 suppression **(Fig. 3I: Left)**, whereas G12R-RR cells leverage EGFR to drive RAS^WT^-GTP loading through a signaling axis largely insensitive to daraxonrasib/CypA engagement **(Fig. 3I: Right)**.

### Cell Lines Resistant to Daraxonrasib Recapitulate Stoichiometric Adaptations

Standard phenotypic assays confirmed that -RR derivative lines were insensitive to increasing daraxonrasib concentrations by MTT and colony formation assays **(Fig. 4A-B)** and maintained ERK phosphorylation at concentrations that fully suppressed signaling in parental cells **(Fig. 4C)**, confirming stable MAPK pathway reactivation. Whole exome sequencing of the resistant lines showed no acquired co-alterations in the RAS network (Methods).

**Figure 4.**
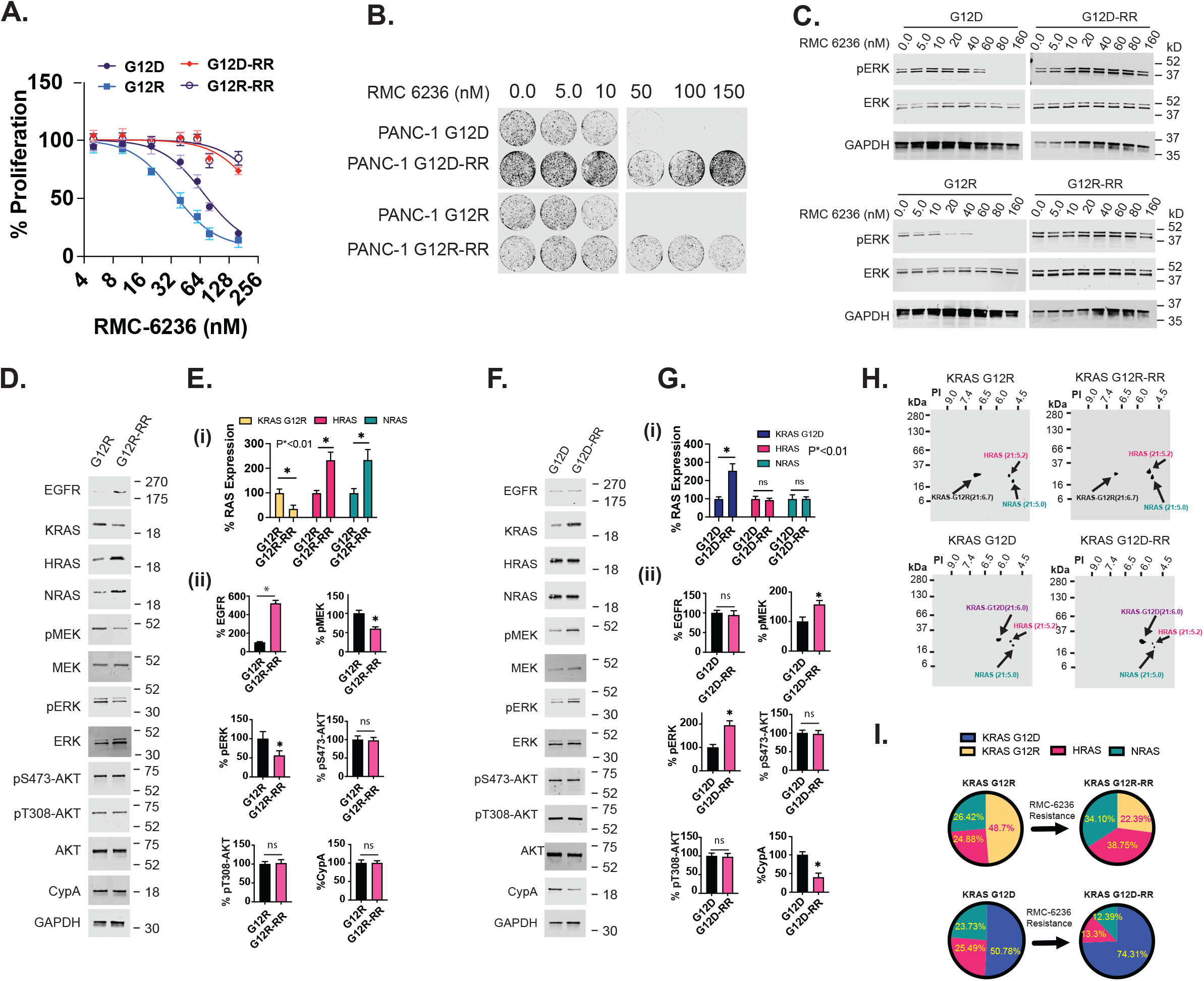
Distinct Stoichiometric Adaptations Confer Daraxonrasib Resistance in KRAS^G12R^ and KRAS^G12D^ PDAC Cells. **A**. PANC-1 G12D and G12R cells were treated with their IC^50^ concentrations of RMC-6236 for greater than 10 passages to develop RMC-6236 resistant lines (-RR). Both PANC-1 KRAS^G12D^-RR and G12R-RR cells did not show any measurable change in IC^50^ by MTT. **B**. Colony formation assay of PANC-1 G12D, G12D-RR, G12R, and G12R-RR cells treated with increasing concentrations of RMC-6236 for 14 days. **C**. Western blot analysis of ERK phosphorylation in response to increasing concentrations of RMC-6236 following 24-hour treatment. **D**. Western blot analysis of RAS/MAPK signaling proteins, CypA, and EGFR in PANC-1 G12R and G12R-RR cells. **E**. Quantification of protein and phosphoprotein expression from panel D. Data represent mean ± SD from three independent experiments. **F**. Western blot analysis of RAS/MAPK signaling proteins, CypA, and EGFR in PANC-1 G12D and G12D-RR cells. **G**. Quantification of protein and phosphoprotein expression from panel F. Data represent mean ± SD from three independent experiments. **H**. Two-dimensional gel electrophoresis and western blot analysis of PANC-1 G12R, G12R-RR, G12D, and G12D-RR lysates probed with pan-RAS antibody. KRAS mutant, HRAS, and NRAS isoforms were identified by mass-to-charge ratio (M:C). **I**. Quantification of RAS isoform expression in parental and RMC-6236-resistant cell lines by 2D electrophoresis. Data represent mean from three independent experiments. Statistical differences were calculated by two-tailed unpaired t-test and P values are indicated (for two groups). For experiments with three or more groups statistical comparisons were calculated by one-way ANOVA followed by post hoc Tukey’s test for multiple comparisons, and P values are indicated in the panels. Each experiment is representative of three experiments; quantifications represent mean and error bars represent standard deviations from three separate experiments

PANC-1 G12R-RR cells showed marked EGFR, HRAS, and NRAS upregulation with modest pMEK/pERK reductions relative to parental cells **(Fig. 4D-E)**. Conversely, PANC-1 G12D-RR cells showed significantly increased KRAS^G12D^ expression correlating with elevated pMEK/pERK; EGFR, HRAS, and NRAS expression remained stable **(Fig. 4F-G)**, and CypA decreased relative to parental cells **(Fig. 4F-G)**. pAKT was unchanged in all cells, indicating MAPK-specific reactivation rather than PI3K/AKT engagement.

Two-dimensional gel electrophoresis quantifying RAS isoform redistribution **(Fig. 4H)** confirmed that G12R-RR cells upregulated HRAS/NRAS while downregulating KRAS^G12R^, shifting stoichiometry from KRAS^MUT^-to RAS^WT^-dominant **(Fig. 4I: Top)**, whereas G12D-RR cells increased KRAS^G12D^ without substantial RAS^WT^ change, retaining a KRAS^MUT^-dominant stoichiometry **(Fig. 4I: Bottom)**, consistent with previously reported RAS isoform stoichiometry boundaries (26–28).

To validate these findings across additional models, we examined KRAS^G12D^ lines (ASPC-1, SW1990, Pa-14C) and KRAS^G12R^ lines (TccPan-2, PSN1), generating -RR derivatives by continuous low-dose treatment for 6 weeks. KRAS^G12D^-RR lines showed modest KRAS decreases relative to controls, opposite the PANC-1 G12D pattern, indicating KRAS^MUT^ upregulation is not obligatory for resistance; pMEK/pERK still rebounded above 4-hour treatment levels with sustained RAS^WT^-GTP loading **(Fig. S3B: Left)**. *KRAS*G12R lines showed elevated basal EGFR/pEGFR under standard culture, discordant with the PANC-1 G12R cells (13); KRAS^G12R^-RR cells did not further upregulate EGFR but instead showed compensatory RAS^WT^ upregulation, as in PANC-1 G12R cells **(Fig. S3B: Right)**.

To assess RAS^WT^ biochemical activity in resistant cells, RAS-GTP pulldown assays showed that daraxonrasib reduced KRAS^G12R^-GTP binding to RAF1-RBD in G12R-RR cells, but RAS^WT^-GTP remained RBD-bound despite drug presence **(Fig. 5A: i)**, contrasting with sensitive parental lines **(Fig. 2A)**. Parallel CypA-GST pulldown confirmed that the daraxonrasib/CypA complex selectively bound KRAS^G12R^ but had limited engagement with RAS^WT^ **(Fig. 5A: ii)**, demonstrating that RAS^WT^-GTP remains free and capable of effector binding in resistant cells, a possible biochemical bypass mechanism explaining how G12R-RR cells recover MAPK activity despite effective KRAS^G12R^ inhibition.

**Figure 5.**
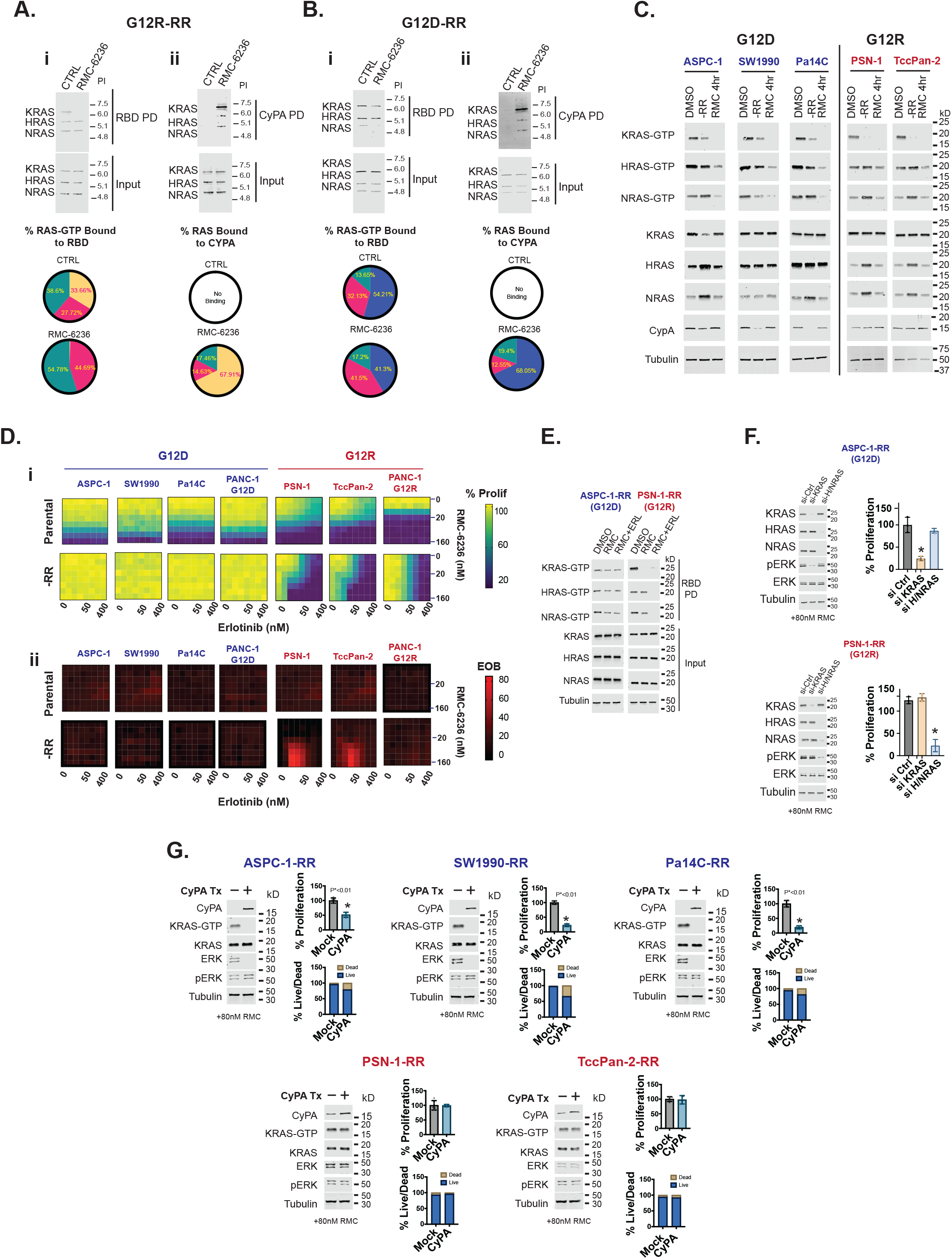
RAS pathway reactivation mechanisms under sustained daraxonrasib treatment. **A**. RBD-IEF analysis of PANC-1 G12R-RR cells treated with vehicle (DMSO) or 80 nM RMC-6236 for 24 hours, immunoblotted with pan-RAS antibody (i:top). Quantification of RAS-GTP isoforms bound to RBD in vehicle (DMSO) versus RMC-6236-treated treatment (i:bottom). CypA-GST/RMC-6236 pulldown of RAS-GTP from PANC-1 G12R-RR cells (ii:top). Quantification of RAS-GTP isoforms bound to the CypA/RMC-6236 complex (ii; bottom). **B**. RBD-IEF analysis of PANC-1 G12D-RR cells treated with vehicle (DMSO) or 80 nM RMC-6236 for 24 hours, immunoblotted with pan-RAS antibody (i:top). Quantification of RAS-GTP isoforms bound to RBD in vehicle (DMSO) versus RMC-6236-treated treatment (i:bottom). CypA-GST/RMC-6236 pulldown of RAS-GTP from PANC-1 G12D-RR cells (ii:top). Quantification of RAS-GTP isoforms bound to the CypA/RMC-6236 complex (ii; bottom). **C**. Cell lines with diverse genetic backgrounds harboring KRAS^G12D^ or G12R mutations were treated with either vehicle (DMSO) or 40 nM RMC-6236 for 4 h or 4 weeks (the latter timepoint corresponding to cells that had regained proliferation). Cells were subjected to active RAS RBD pulldown, and pulldown and input lysates were resolved by SDS-PAGE and probed for K-, H-, and N-RAS; input lysates were additionally probed for CypA and tubulin as loading controls. **D**. (i)Cell lines harboring KRAS^G12D^ or G12R mutations were treated with RMC-6236, erlotinib, or the combination at increasing doses, and proliferation (% prolif.) was measured by CellTiter-Glo assay. (ii) Excess over Bliss was calculated from proliferation data to identify conditions of synergy. **E**. Two representative -RR cell lines harboring KRAS^G12D^ or KRAS^G12R^ were treated with 80 nM RMC-6236 alone or in combination with 50 nM erlotinib, and active RAS RBD pulldown assays were performed; input and pulldown fractions were resolved by SDS-PAGE and probed for K-, H-, and N-RAS. **F**. ASPC-1-RR and PSN-1-RR cells, which are maintained in RMC-6236-containing media, were transfected with control siRNA, KRAS siRNA, or HRAS+NRAS siRNA. Twenty-four hours later, cells were lysed, and lysates were resolved by SDS-PAGE and probed for H-, N-, and K-RAS. At 72 h, cell proliferation was measured by MTT assay. **G**. Representative RMC-6236-resistant (-RR) lines were maintained in RMC-6236-containing media and transfected with a pcDNA expression vector for CypA. Cells were subjected to active RAS RBD pulldown; the pulldown fraction was resolved by SDS-PAGE and probed for KRAS (KRAS-GTP band). Input was resolved by SDS-PAGE and probed for CypA, KRAS, total ERK, pERK, and tubulin. Transfected cells were also compared to empty vector control by MTT assay, and live/dead staining assay 72 hrs post transfection. All experiments were three times. Statistical differences were calculated by two-tailed unpaired t-test and P values are indicated (for two groups). For experiments with three or more groups statistical comparisons were calculated by one-way ANOVA followed by post hoc Tukey’s test for multiple comparisons, and P values are indicated in the panels. Each experiment is representative of three experiments; quantifications represent mean and error bars represent standard deviations from three separate experiments.

In G12D-RR cells, daraxonrasib caused only modest reductions in both KRAS^G12D^ and RAS^WT^ RBD binding **(Fig. 5B: i)**, contrasting with the complete RBD-binding loss seen in parental cells **(Fig. 2A)**. CypA-GST pulldown showed the daraxonrasib/CypA complex preferentially bound KRAS^G12D^ with negligible RAS^WT^ interaction **(Fig. 5B: ii)**.

To compare the isogenic system with cell lines of distinct genetic backgrounds, we performed RAS activity assays across all lines. In *KRAS*G12D lines, 4-hour daraxonrasib treatment ablated KRAS-GTP signal and reduced HRAS/NRAS-GTP, consistent with transactivation sustaining RAS^WT^-GTP (13,14,16,17); in -RR derivatives, both KRAS^MUT^ and RAS^WT^ showed increased RBD binding relative to 4-hour treatment, correlating with elevated RAS^WT^ expression and CypA downregulation **(Fig. 5C)**. KRAS^G12R^ lines instead showed sustained KRAS^MUT^-GTP ablation with no rescue, while HRAS/NRAS-GTP increased in -RR cells, mirroring PANC-1 G12R behavior. Notably, only *KRAS*G12D lines showed CypA downregulation and partial KRAS^MUT^-GTP rebound, while only *KRAS*G12R lines showed consistent RAS^WT^-GTP upregulation and EGFR expression **(Fig. 5C)**.

PSN-1 and TccPan-2 (*KRAS*G12R) cells robustly express EGFR at baseline **(Fig. S3B)**, a feature not observed in the PANC-1 isogenic cell lines. Given that KRAS^G12R^ is an intrinsically weaker oncogenic driver with limited ERK/MAPK capacity despite higher CRAF affinity (13,20,21), we postulate EGFR overexpression here reflects compensatory adaptation for tumor growth and survival (13). To define the functional role of EGFR in daraxonrasib sensitivity and resistance, we employed combinatorial drug screening.

Combination treatment with daraxonrasib and erlotinib showed: *KRAS*G12D parental lines were daraxonrasib-sensitive but required higher doses than *KRAS*G12R lines and were unaffected by erlotinib, consistent with lacking EGFR **(Fig. 5D: i)**; KRAS^G12D^-RR derivatives were fully resistant to both agents and their combination, indicating EGFR-independent ERK reactivation. Among *KRAS*G12R lines, sensitivity tracked with baseline EGFR: EGFR-high PSN-1/TccPan-2 parental cells were daraxonrasib-sensitive and partially erlotinib-sensitive, while EGFR-low PANC-1 G12R cells were daraxonrasib-hypersensitive but erlotinib-insensitive **(Fig. 5D: i)**. After resistance, PSN-1-RR and TccPan-2-RR lost daraxonrasib sensitivity but gained erlotinib sensitivity, with strong combination synergy **(Fig. 5D: i-ii)**. PANC-1 G12R-RR cells instead became erlotinib-hypersensitive, remained daraxonrasib-resistant, and showed no synergy, associated with less KRAS^G12R^ and increased HRAS/NRAS-GTP **(Fig. 4D and 5A: i)**. Furthermore, daraxonrasib alone or in combination with erlotinib had a negligible effect on KRAS signaling in ASPC-1-RR cells **(Fig. 5E: Left)**, consistent with the proliferation data in **Fig. 5D: i**. In PSN-1-RR cells, daraxonrasib alone decreased RBD-bound KRAS^G12R^-GTP while RAS^WT^-GTP RBD binding remained intact; the combination of daraxonrasib with erlotinib, however, robustly decreased all three RAS-GTP isoforms **(Fig. 5E)**.

To confirm on-target resistance, siRNA knockdown of KRAS or HRAS/NRAS in -RR cells (daraxonrasib-containing media) showed that KRAS knockdown markedly reduced ASPC-1-RR proliferation with negligible HRAS/NRAS effect, while the converse held in PSN-1-RR cells **(Fig. 5F)**. These results infer that pathway reactivation and resistance arise from continued signaling by drug-free RAS isoforms, with the EGFR resistance mechanism dependent on wild-type RAS.

Since G12D-RR cells downregulated CypA expression **(Fig. 4F)** concurrent with restored KRAS^G12D^-GTP **(Fig. 2A and Fig. 5B-C)**, we hypothesized that reduced CypA availability drives resistance in this context. Comparing PANC-1 KRAS^G12D^ and G12D-RR cells, we observed that CypA overexpression resulted in loss of KRAS-GTP in G12D-RR cells following RMC-6236 treatment, whereas RAS-GTP isoform changes did not differentiate between mock and CypA in the parental lines **(Fig. S3C)**. In daraxonrasib-containing growth media, restoring CypA expression in non-isogenic G12D-RR cells sharply reduced KRAS-GTP, pERK, and proliferation **(Fig. 5G: Top)**, whereas CypA overexpression had no effect in G12R-RR cells, which retain endogenous CypA **(Fig. 5G: Bottom)**, indicating decreased CypA expression as a possible route to sustained KRAS^G12D^-GTP in resistant G12D cells.

### KRAS^G12R^ Patient Achieves Prolonged Response to Daraxonrasib Before Developing Acquired Resistance

In March 2025, a 69-year-old man with metastatic PDAC presented after five prior lines of therapy **(Fig. 6A)**, having been diagnosed in May 2022 with *KRAS*G12R-mutant disease. First-line FOLFIRINOX (June-November 2022) produced a CA19-9 and radiographic response; due to chemotherapy-induced neuropathy, treatment was de-escalated to FOLFIRI maintenance until progression in March 2023, yielding a combined PFS of 9.7 months, exceeding the 6.8-month median reported for unselected PDAC patients on FOLFIRINOX (29), consistent with *KRAS*G12R conferring modestly improved outcomes (20) **(Fig. 6A: i)**. Second-line gemcitabine/nab-paclitaxel (April 2023) produced response and stability but was discontinued after 2.8 months for worsening neuropathy. Third-line daraxonrasib (August 2023) yielded robust initial responses consistent with KRAS^G12R^ hypersensitivity, though toxicities required dose reduction from 400 to 200 mg daily; the patient progressed in June 2024, approximately 10 months after initiation **(Fig. 6A: ii)**. Fourth- and fifth-line experimental antibody-drug conjugate (ADC) therapies (targeting c-MET and claudin 18.2) produced rapid progression after 11 and 5 weeks, respectively. The patient achieved nearly 24 months of disease control across the first three lines (7), and Tempus xT profiling after daraxonrasib resistance showed no acquired alterations from diagnosis **(Fig. 6A: iii)**.

**Figure 6.**
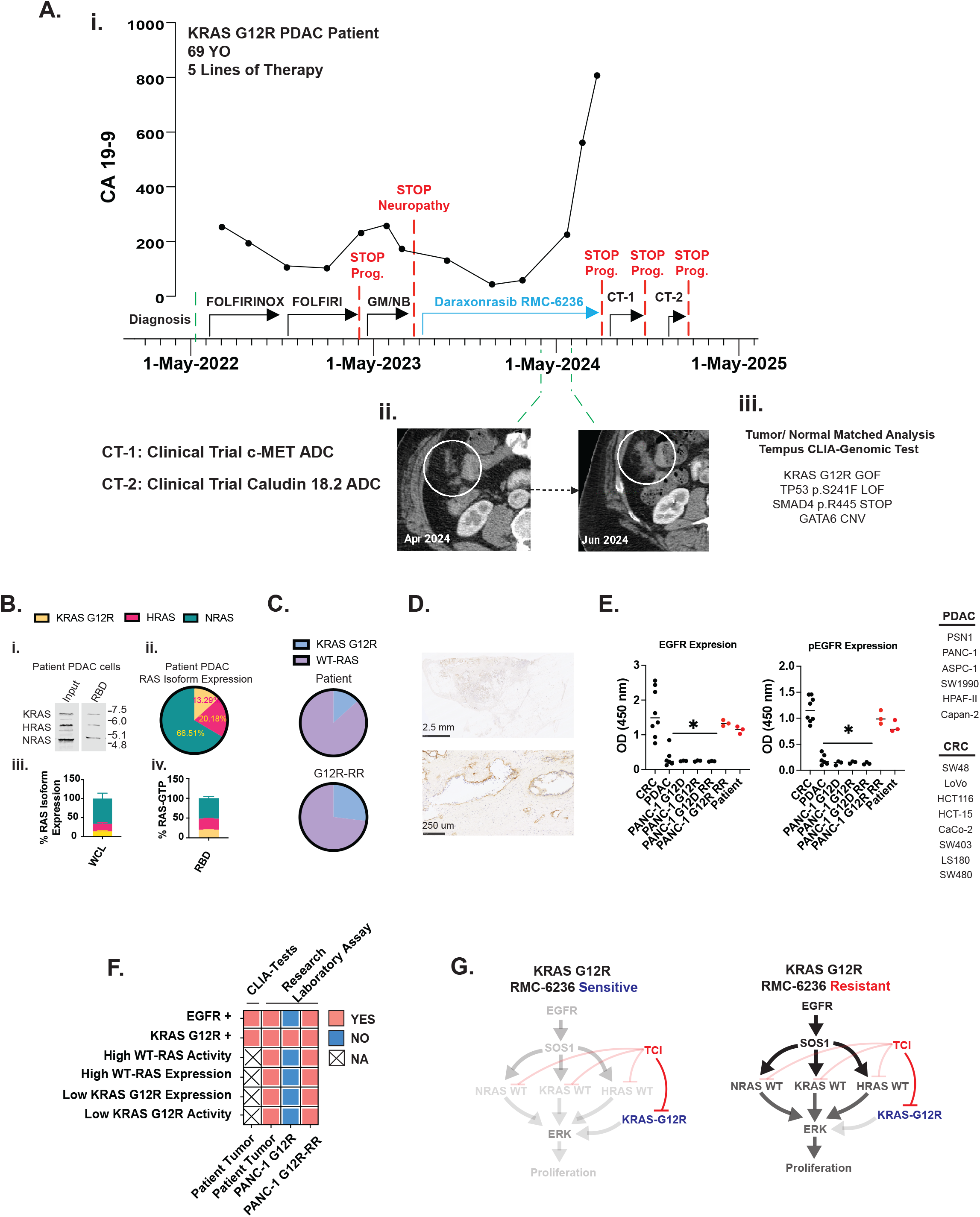
Molecular Profiling of Daraxonrasib-Resistant KRAS^G12R^ Patient Tumor Validates Preclinical Resistance Mechanisms. **A. (i)** Treatment timeline versus carbohydrate antigen 19-9 (CA19-9) levels; showing lines of therapy received before presentation to the LaBahn Pancreatic Cancer Program. Each therapy was discontinued (red dashed lines) due to disease progression or chemotherapy-induced peripheral neuropathy (CIPN). Third-line daraxonrasib (RMC-6236) was initiated in August 2023. Despite initial response, the patient subsequently experienced rising CA19-9 levels indicating disease progression on daraxonrasib. **(ii)** Serial computed tomography (CT) imaging demonstrating a representative peritoneal metastatic nodule with progressive enlargement from April 2024 to June 2024 during daraxonrasib treatment, confirming radiographic progression. Following daraxonrasib discontinuation, the patient enrolled in two sequential clinical trials (fourth-line antibody-drug conjugate targeting c-MET; fifth-line antibody-drug conjugate targeting claudin 18.2), both resulting in rapid disease progression. **(iii)** Tempus xT next-generation sequencing performed after progression on fifth-line therapy revealed retained KRAS^G12R^ mutation with no new genomic alterations compared to pre-treatment molecular profiling. **B**. Viable tumor cells were isolated from a patient peritoneal metastatic lesion by fluorescence-activated cell sorting (FACS) following biopsy. Active RAS-GTP was captured by RBD pulldown, and both input lysates and RBD-bound material were analyzed by one-dimensional isoelectric focusing (1D-IEF) with pan-RAS immunoblotting **(i)**. Quantification of RAS isoform stoichiometry from three samples showing total RAS isoform distribution (**ii, iii)**, and RAS-GTP isoform composition **(iv)**. **C**. Comparison of KRAS^MUT^, HRAS^WT^, and NRAS^WT^ stoichiometric distribution between laboratory-generated daraxonrasib-resistant cell lines (PANC-1 G12R-RR) and patient-derived tumor cells (KRAS^G12R^) with acquired daraxonrasib resistance. **D**. Immunohistochemical analysis of the tumor biopsy specimen performed by CLIA-certified pathology services (Wisconsin Diagnostic Labs) confirmed EGFR positivity. **E**. ELISA assays for EGFR (left) and phosphorylated EGFR (right) for cell line libraries, PANC-1 isogenic cells and their RMC-6236 resistant derivatives (-RR) and patient cells from the liver metastatic site. **F**. Summary of molecular profiling from the patient’s metastatic liver lesion compared to laboratory-generated cell line resistance models, using FDA-approved CLIA-certified testing for actionable biomarkers on serial biopsies and non-CLIA research assays. **G**. Biochemical analysis of PANC-1 isogenic cell lines (KRAS^G12D^ and G12R) and their RMC-6236-resistant clones demonstrated that ERK pathway reactivation through EGFR and wild-type H-RAS/N-RAS signaling mediates resistance to RMC-6236. Statistical differences were calculated by one-way ANOVA followed by post hoc Tukey’s test for multiple comparisons, and P values are indicated in panel E. Each experiment is representative of three experiments; quantifications represent mean and error bars represent standard deviations from three separate experiments.

A resected metastatic lesion was sorted by FACS for viable cancer cells and analyzed by RBD-IEF, revealing markedly elevated RAS^WT^ expression and activation with low KRAS^G12R^ levels **(Fig. 6B)**. The patient’s tumor showed a more pronounced RAS^WT^-dominant stoichiometry (86.67% RAS^WT^) than laboratory-derived KRAS^G12R^-RR lines (72.85% RAS^WT^) **(Fig. 6C)**, recapitulating our preclinical resistance models with an even greater reliance on RAS^WT^ signaling.

Given that isogenic G12R-RR cells acquire EGFR expression driving RAS^WT^ activity preclinically, CLIA-certified IHC confirmed EGFR positivity in the patient’s tumor **(Fig. 6D)**. ELISA quantifying total and phosphorylated EGFR across CRC lines (positive control), treatment-naive PDAC lines (negative control), parental and resistant isogenic cells, and patient-derived cells showed that both G12R-RR cells and patient tumor cells had comparable EGFR and pEGFR levels, substantially exceeding treatment-naive PDAC and approaching CRC levels **(Fig. 6E)**, demonstrating convergence toward an EGFR-dependent, RAS^WT^-reliant signature in both laboratory and clinical contexts **(Fig. 6F)**.

Since the daraxonrasib/CypA complex preferentially binds KRAS^MUT^-GTP over RAS^WT^-GTP **(Fig. 2A-D)**, and KRAS^G12R^ cannot transactivate RAS^WT^ **(Fig. 1C-D)**, these data explain the hypersensitivity of treatment-naive *KRAS*G12R PDAC to daraxonrasib **(Fig. 6G, Left)**. Acquired/inherent EGFR expression then allows resistant cells to activate RAS^WT^, bypassing daraxonrasib blockade and restoring MAPK flux despite continued KRAS^G12R^ inhibition **(Fig. 6G, Right)**. Given the rapid progression on fourth- and fifth-line therapies (11 and 5 weeks) and the strong concordance between patient and G12R-RR signaling profiles **(Fig. 6F)**, we hypothesized that our resistance models could prospectively guide salvage therapy selection.

### Preclinical Modeling and Rational Drug Selection for Disease Control in Late-Line Daraxonrasib-Resistant PDAC

As traditional organoid culture from this heavily pretreated patient’s biopsy was unsuccessful, we developed a rapid 3D bioprinting platform (Greiner Bio-One) generating toroidal structures in ultra-low attachment conditions, enabling accelerated drug-sensitivity assessment via structural integrity and anoikis resistance (30–34) **(Fig. S4)**.

We first validated this approach using PANC-1 isogenic lines and their -RR derivatives. G12D cells showed toroid disruption at 60 nM daraxonrasib, while G12D-RR toroids remained intact at all concentrations **(Fig. 7A: Left)**; G12R cells dissolved at 20 nM while G12R-RR toroids remained intact **(Fig. 7A: Right)**. Cosine similarity quantification of structural integrity detected this dissolution **(Fig. S6A)** and was orthogonally validated by trypan blue viability **(Fig. S6B)** and CellTiter-Glo luminescence **(Fig. S6C)**, with significant reductions in both readouts at wells showing toroid loss. Cosine similarity showed greater sensitivity than trypan-blue-exclusion and agreed more closely with CellTiter-Glo, consistent with daraxonrasib acting predominantly as cytostatic rather than cytotoxic at these concentrations. All three readouts showed significant concordance (Pearson correlation, R^2^ > 0.8), supporting toroid integrity as a reliable surrogate for anti-tumor activity **(Fig. S6G)**.

**Figure 7.**
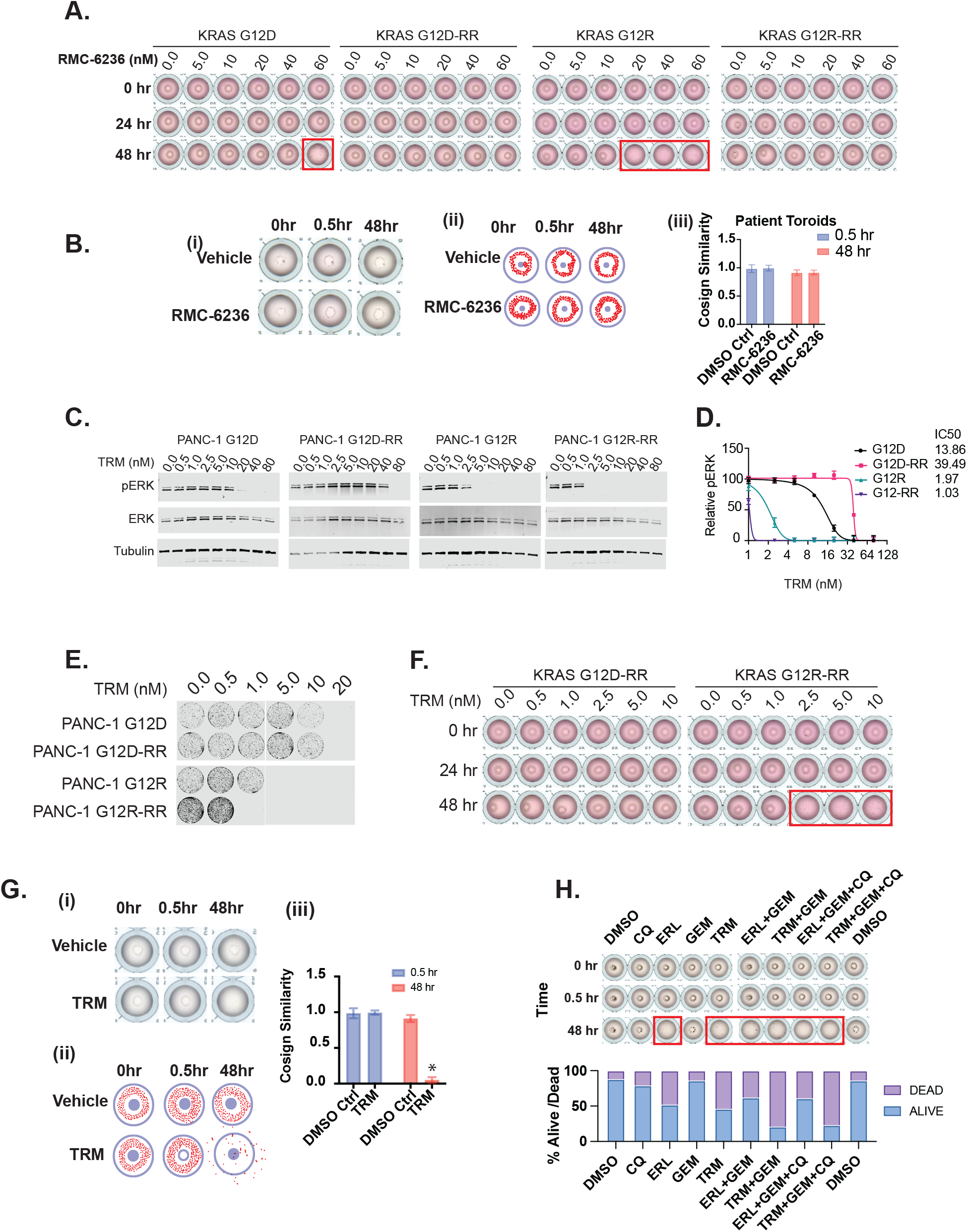
Preclinical Modeling Identifies Trametinib Sensitivity in Daraxonrasib-Resistant KRAS^G12R^ PDAC. A. PANC-1 isogenic cell lines (G12D, G12R) and their daraxonrasib (RMC-6236)-resistant derivatives (G12D-RR, G12R-RR) were bioprinted into three-dimensional toroidal structures under ultra-low attachment conditions and treated with increasing concentrations of RMC-6236 in complete medium for 48 hours. Toroidal disruption (red boxes) indicates drug sensitivity and loss of anoikis resistance. B. Patient-derived tumor cells were bioprinted into three-dimensional toroids under ultra-low attachment conditions in organoid growth medium and treated with 40 nM RMC-6236. **(i)** Representative images at 30 minutes and 48 hours post-treatment. **(ii)** Pixel mapping analysis within spherical markers to quantify structural deviation from baseline toroidal geometry. **(iii)** Cosine similarity scores compared to 0-hour timepoint demonstrate maintained structural integrity over 48 hours of treatment, indicating persistent daraxonrasib resistance. C. PANC-1 isogenic cells and their RMC-6236-resistant derivatives were treated with increasing concentrations of trametinib for 24 hours, cell lysates were prepared and resolved by DS-PAGE and blotted for phosphorylated ERK (pERK), ERK, and Tubulin. D. Quantification of pERK/ERK fluorescence from three separate experiments for PANC-1 isogenic cells and their RMC-6236-resistant derivatives from **panel C**. E. PANC-1 G12R and G12D isogenic lines along with their RMC-6236 resistance (-RR) clones were plated for colony formation and treated with increasing concentrations of trametinib for 2 weeks. F. PANC-1 RMC-6236-resistant derivatives (G12D-RR, G12R-RR) were bioprinted into three-dimensional toroidal structures under ultra-low attachment conditions and treated with increasing concentrations of trametinib (TRM) in complete medium for 48 hours. Toroidal disruption (red boxes) indicates drug sensitivity and loss of anoikis resistance. G. Patient-derived tumor cells were bioprinted into three-dimensional toroids under ultra-low attachment conditions in organoid growth medium and treated with 5nM trametinib (TRM). **(i)** Representative images at 30 minutes and 48 hours post-treatment. **(ii)** Pixel mapping analysis within spherical markers to quantify structural deviation from baseline toroidal geometry. **(iii)** Cosine similarity scores compared to 0-hour timepoint demonstrate loss of structural integrity over 48 hours of treatment, indicating that the daraxonrasib-resistant patient cells are hyper-sensitive to low dose MEKi. H. Patient-derived tumor cells were bioprinted into three-dimensional toroids under ultra-low attachment conditions in organoid growth medium. Toroids were treated with either DMSO (vehicle), chloroquine (CQ; 40 µM), erlotinib (ERL; 500 nM), gemcitabine (GEM; 20 µM), trametinib (TRM; 5 nM), or the indicated combinations. Red boxes indicate toroid dissolution. Following toroid imaging, a trypan blue exclusion assay was performed to determine the percentage of viable and dead cells after treatment with individual drugs and combinations.

Despite limited viable tumor cells from the patient biopsy, we generated patient-derived toroids using this platform. Treatment with 40 nM daraxonrasib for 48 hours showed maintained structural integrity with no morphological change **(Fig. 7B)**, recapitulating the resistant phenotype of PANC-1 G12R-RR cells and confirming persistent resistance in the patient’s tumor.

The resistance signaling network suggested two targeted therapies to include in a combination treatment approach: EGFR inhibition or MEK inhibition with trametinib. Testing increasing doses in isogenic lines, KRAS^G12R^-RR cells were significantly more trametinib-sensitive than all other lines, which remained resistant **(Fig. S5)**.

We further investigated MEK inhibition given KRAS^G12R^’s weaker signaling and reduced MEK phosphorylation. Trametinib preferentially binds unphosphorylated MEK, blocking RAF-mediated activation (35,36), suggesting daraxonrasib-resistant KRAS^G12R^ PDAC, with its KRAS^G12R^-to-RAS^WT^ signaling shift and reduced pMEK/pERK **(Fig. 4D)**, might show increased trametinib sensitivity. Treating isogenic models and resistant derivatives confirmed this: PANC-1 G12D and G12D-RR were trametinib-resistant, while PANC-1 G12R and G12R-RR were sensitive, with G12R-RR showing greater sensitivity than parental G12R cells **(Fig. 7C-D)**, correlating with IC50 values from proliferation **(Fig. S5)** and colony formation assays **(Fig. 7E)**, confirming trametinib suppresses MAPK signaling in *KRAS*G12R contexts regardless of daraxonrasib resistance status (13).

Trametinib sensitivity was further evaluated in rapid-bioprinted toroids from G12D-RR and G12R-RR cells: PANC-1 G12D-RR toroids remained intact across all concentrations, while G12R-RR toroids disrupted at concentrations as low as 2.5 nM **(Fig. 7F)**, confirming allele-specific MEK-inhibitor vulnerability. Cosine similarity, trypan blue, and CellTiter-Glo analyses on corresponding wells were concordant with this toroid dissolution data **(Fig. S6F-G)**.

Patient-derived toroids treated with vehicle or low-dose (5 nM) trametinib for 48 hours showed complete disruption with trametinib **(Fig. 7G)**, indicating tumor hypersensitivity to MEK inhibition comparable to PANC-1 G12R-RR cells and supporting trametinib-based therapy. Combination regimens were prioritized over monotherapy per our institutional molecular tumor board guidelines. Residual tumor tissue was bioprinted into toroids to screen combinations of FDA-approved agents available for off-label use under our protocol: chloroquine (the active metabolite of hydroxychloroquine), erlotinib, trametinib, and gemcitabine. Due to limited tissue, drug concentrations were selected to approximate reported intratumoral concentrations in pancreatic cancer patients (13,22,37,38). We observed that trametinib and erlotinib, alone or in combination with other drugs, caused dissociation of toroid structures; however, the greatest degree of cell killing was observed when trametinib was paired with gemcitabine **(Fig. 7H)**.

### Combination Therapy Regimen Achieves Disease Control After Daraxonrasib Resistance

Given overlapping erlotinib/trametinib toxicity profiles (rash, diarrhea) and tolerability concerns in this heavily pretreated patient, we selected single-agent trametinib over combination EGFR/MEK inhibition, based on: (i) the modest benefit of erlotinib plus gemcitabine over gemcitabine alone in first-line non-metastatic PDAC (PFS 6.2 vs. 5.9 months) (39); (ii) superior 6.8-month median PFS with trametinib-based therapy in early-line *KRAS*G12R PDAC (13); and (iii) case report evidence supporting gemcitabine plus MEK inhibition in late-line *KRAS*G12R PDAC despite prior gemcitabine exposure (40). At presentation, after 5 prior lines, CT showed progressive hepatic, pulmonary, and peritoneal disease versus a scan 6-7 weeks earlier. Since MEK inhibition is likely primarily cytostatic **(Fig. S6)**, gemcitabine was added to target the proliferative escape fraction, given tumor heterogeneity in EGFR/RAS^WT^ dependence; hydroxychloroquine was added based on our prior data showing an additive trametinib effect via suppressed autophagy-mediated resistance (13,22,41).

The patient started trametinib (1 mg daily; approved dose 2 mg), hydroxychloroquine (600 mg twice daily), and gemcitabine (800 mg/m^2^ days 1, 8, 15 of a 28-day schedule; approved dose 1000 mg/m^2^) in March 2025 **(Fig. 8A)**, achieving a robust CA19-9 response and 5 months of imaging stability **(Fig. 8B)**. Treatment was well tolerated, with only mild skin rash, mouth sores, fatigue, and cytopenia. After approximately 5 months, the patient developed aggressive progression (representative liver lesion, **Fig. 8C**), worsening pain, and rapid clinical deterioration, succumbing in late August 2025 with an overall survival of 40 months from diagnosis.

**Figure 8.**
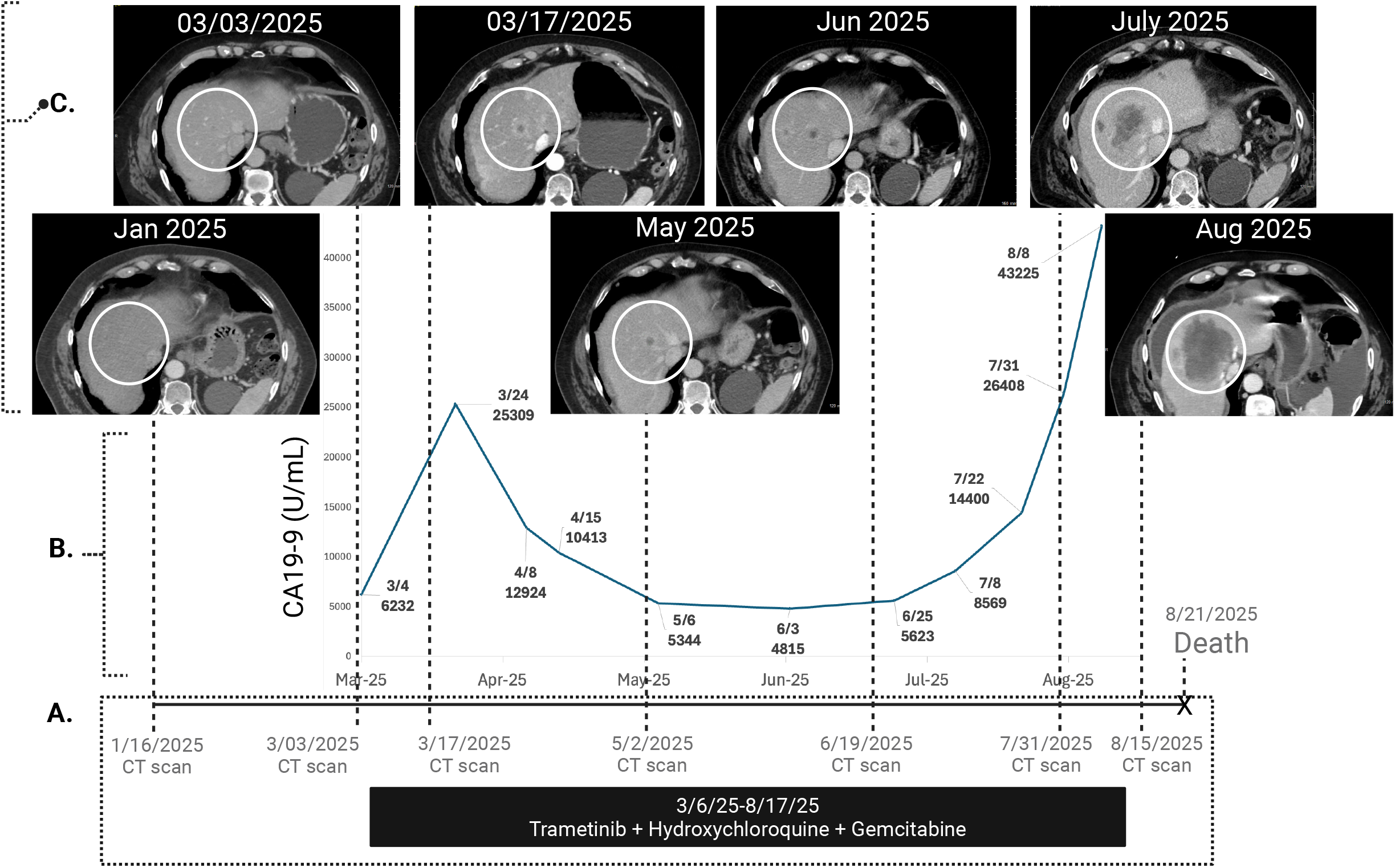
Sixth-Line Therapy Achieves Durable Response in Daraxonrasib-Resistant Metastatic PDAC. Patient treatment and outcome**;(A)** Represents the timeline; **(B)** Represents the serial levels of cancer antigen 19-9 (CA19-9) while on the MEK inhibitor-based combination therapy; **(C)** Shows serial computed tomography (CT) scans. A representative lesion in the liver, marked by the white circle, is followed over time. As presented, within the first 2-3 weeks of treatment initiation patient had fast progression of disease. This is represented by emergence of a new liver lesion on the 3/17/25 imaging, which was absent on 3/3/25 CT scan, and a significant rise in CA19-9 from 6,236 in 3/4/25 to 25,309 on 3/24. From 3/24, patient had response to therapy represented by the decrease in CA19-9 and achievement of stability of the represented liver lesion. From July, patient experienced rise in CA19-9, and by 7/31 CT showed significant progression (4.8 months from treatment initiation), with a rapidly growing lesion, nearly doubling in size form 7/31/25 to 8/15/25.

## DISCUSSION

This patient’s trajectory underscores three insights: KRAS allele identity shapes resistance mechanisms; real-time translational research can bridge the gap between investigational agent failure and salvage therapy; and molecularly informed adaptive strategies can extend survival even in heavily pretreated advanced disease. KRAS^G12R^’s inability to transactivate RAS^WT^ confers both initial hypersensitivity to RAS inhibition and constrained resistance routes, with the potential of creating exploitable vulnerabilities even after progression. The convergence of patient-derived and preclinical resistance mechanisms correlated with rational sixth-line salvage therapy in a setting lacking evidence-based guidelines.

*KRAS*G12R has distinct biophysical properties relative to the more common *KRAS*G12D and *KRAS*G12V variants: it lacks SOS1 allosteric activation and cannot serve as an NF1 stoichiometric sink, attenuating RAS^WT^ activity (13– 17). This reduced holistic RAS pathway activity likely explains decreased tumorigenic potential in murine models (13,20,22), consistent with *KRAS*G12R’s clinical enrichment in well-to moderately-differentiated localized PDAC versus *KRAS*G12D’s enrichment in metastatic disease (20,42).

Many patients who develop targeted therapy resistance, including to KRAS^G12C^ inhibitors like sotorasib and adagrasib, lack identifiable genomic co-alterations (10), reflecting protein-level signaling remodeling beyond sequencing’s reach. While next-generation sequencing remains essential for resistance profiling, protein expression and signaling adaptation require complementary approaches. Here, no new actionable mutations emerged after daraxonrasib progression; resistance instead likely arose through EGFR and RAS^WT^ protein-level upregulation and activation. Our integrated functional approach, combining direct intraoperative RAS activity measurement, RAS network stoichiometry assessment, and *ex vivo* drug sensitivity testing on surgically obtained tissue, identified the EGFR/RAS^WT^ axis as the operative mechanism, underscoring the value of pairing protein-level studies and functional assays with genomic profiling in targeted therapy resistance research.

KRAS^G12D^ PDAC has been reported resistant to combined MEK and autophagy inhibition with gemcitabine (43), whereas KRAS^G12R^ tumors are relatively sensitive, with an extraordinary case achieving 22 months of disease control (13), attributed to the lower basal pMEK/ERK threshold characteristic of KRAS^G12R^ tumors (21), favoring trametinib’s selective binding to unphosphorylated MEK (13). Our KRAS^G12R^ models suggested EGFR/PI3K/AKT resistance routes were unlikely, with reactivation instead driven by an EGFR/RAS^WT^-driven ERK program. Given our patient’s RAS^WT^-dominant signaling profile, we hypothesized that trametinib, gemcitabine, and hydroxychloroquine would similarly exert antitumor activity.

Conversely, our daraxonrasib-resistant KRAS^G12D^ models retained mutant KRAS-GTP with CypA downregulation, and MEK inhibition did not alter proliferation, suggesting KRAS^G12D^ PDAC may instead require CypA-independent KRAS inhibitors, phospho-MEK-targeting MEK inhibitors, or ERK inhibitors as they become available.

The absence of published daraxonrasib resistance data at presentation necessitated *de novo* investigation under significant time pressure. We prioritized platforms returning actionable data within days, since organoids are slow to establish and show lower KRAS^G12R^ take rates (13,20,21). We instead used *ex vivo* organotypic slice cultures and patient-derived toroid models, established the day of biopsy and preserving native tumor architecture and heterogeneity, enabling biologically relevant drug sensitivity and signaling assessment without prolonged model generation timelines. Though non-passageable and material-limited, these models provided time-critical, actionable readouts within the clinical action window.

A recent complementary study using allele-anchored CRISPR-Cas9 screens across *KRAS*-mutant PDAC lines identified genetic modulators of KRASi sensitivity, including EGFR, CK2, and YAP, and similarly showed allele-specific resistance landscapes and KRAS dependency (44). Whereas that study used population-level genetic perturbation and high-dose escalation, our work leverages systems biology and patient-derived modeling at long-term low-dose resistance to capture non-genetic, expression-level mechanisms in real time, offering a complementary, clinically actionable framework for allele-informed therapy.

The drivers of EGFR and RAS^WT^ upregulation in resistance remain incompletely defined, though feedback mechanisms offer a plausible explanation: KRAS inhibition relieves transcriptional EGFR suppression, partly via downregulating negative regulators like ERRFI1 (ERBB receptor feedback inhibitor 1), reactivating EGFR-driven MAPK/ERK output through wild-type HRAS and NRAS (45,46), consistent with our KRAS^G12R^-RR models. Supporting RAS^WT^ redundancy as a compensatory mechanism, the pan-KRAS inhibitor BI-2865 shows attenuated efficacy in KRAS^G12R^ lines, reversed by HRAS/NRAS knockdown (47), indicating RAS^WT^ can buffer pan-RAS inhibition. The transcriptional program governing this induction remains unresolved but warrants further investigation.

While EGFR/RAS^WT^ activity was the observed KRAS^G12R^ resistance mechanism, alternative routes remain likely. KRAS^G12R^ amplification is a theoretical possibility, though the biophysical constraints conferring CypA-daraxonrasib hypersensitivity suggest this would require amplification well beyond that seen with KRAS^G12D^. In KRAS^G12D^ contexts, tumors might achieve MEKi-sensitive phenotypes via RAS^WT^-low/RAS^MUT^-low configurations, though such states may be selected against during disease development; this reductionist framework relating RAS^WT^, KRAS^MUT^, and EGFR remains plausible. Our findings do not represent the sole resistance route, but their consistency across preclinical, translational, and clinical response contexts demonstrates this pipeline’s value.

The combination nature of the salvage regimen – trametinib plus hydroxychloroquine plus gemcitabine – precludes attributing disease control to any single agent; the clinical response supports a trametinib-containing strategy rather than direct validation of trametinib sensitivity alone. The choice of this specific combination therapy was based on prior evidence that MEK inhibition is primarily cytostatic (48), motivating gemcitabine’s addition for cytotoxic coverage, and prior data showing an additive hydroxychloroquine-trametinib effect via suppressed autophagy-mediated resistance (13,49).

This case represents an extraordinarily rare outcome in metastatic PDAC, a disease of dismal prognosis with limited late-line options. Our patient achieved approximately 40 months of overall survival with disease control across six lines of therapy, a trajectory virtually unprecedented in the literature. Only half of metastatic PDAC patients are fit for second-line therapy (50), and among these, only 37-48% proceed to third-line treatment (51,52), with median overall survival of 3.5-5.5 months at that stage (53); comparative sixth-line efficacy data are essentially nonexistent. This sustained benefit reflects a multipronged approach of real-time resistance profiling and adaptive therapeutic selection based on the tumor’s adaptations rather than empiric sequencing, enabling repeated disease control as resistance emerged, demonstrating that molecularly informed salvage therapy can achieve meaningful outcomes where evidence-based guidelines do not exist.

## Author Contributions

### Conceptualization

T. McFall, M. Kamgar, G. Hobbs

### Methodology

M. Kamgar, R. Davidson, G.A. Hobbs, D. Dorbin, R. Kurzrock, T. McFall, B.F. Volkman, F.C. Peterson, E. Ke, N. Atallah, F.A. Sayahpour, G. Johnson, J. Yuan

### Investigation

M. Kamgar, R. Davidson, D. Dorbin, G. Scheuber, J. Herrera, Y.D. Seo, J.F. Langenheim, G.A. Hobbs, N.K. Lytle, S. Tsai, R. Kurzrock, T. McFall, P. Jayakrishnan, C. Rajesh, M. Sochor, D.B. Evans, N.K. Chandrashekar, J. Wittmann, M. Ali, E. Ke, N. Atallah, F.A. Sayahpour

### Formal Analysis

MKamgar, R. Davidson, D. Dorbin, G. Scheuber, J. Herrera, Y.D. Seo, J.F. Langenheim, G.A. Hobbs, N.K. Lytle, S. Tsai, R. Kurzrock, T. McFall, P. Jayakrishnan, C. Rajesh, M. Sochor, D.B. Evans, N.K. Chandrashekar, J. Wittmann, M. Ali, A. Szabo

### Resources

B. Evans, S. Tsai, Y.D. Seo, B.F. Volkman, F.C. Peterson, E. Ke, N. Atallah, F.A. Sayahpour

### Writing, Original Draft

J. Herrera, D. Dorbin

### Writing, Review and Editing

M. Kamgar, R. Davidson, N.K. Lytle, S. Tsai, G.A. Hobbs, R. Kurzrock, T. McFall

### Data Curation

M. Aldakkak, R. Davidson, Y.D. Seo, J. Yuan

### Project Administration

R. Davidson, T. McFall

### Supervision

T. McFall, M. Kamgar, G.A. Hobbs

### Funding Acquisition

T. McFall, M. Kamgar, R. Kurzrock, G.A. Hobbs

## Funding

Medical College of Wisconsin, Department of Surgery We Care Fund (T. McFall, M. Kamgar)

Medical College of Wisconsin Cancer Center, Byrne Scholar Fund (T. McFall, R. Kurzrock)

American Cancer Society and Medical College of Wisconsin Cancer Center Institutional Research Grant, IRG #19-138-34 (T. McFall)

National Institutes of Health, National Cancer Institute, 1R01CA276771-01A1 (T. McFall)

National Institutes of Health, L30CA284420, “Utilizing MEK Inhibitors in Precision Medicine Practices for Pancreatic Cancer” (T. McFall)

Pancreatic Cancer Action Network Career Development Award in memory of Skip Viragh, 22-20-HOBB (G.A. Hobbs)

Concern Foundation Career Development Award (G.A. Hobbs)

National Institute of General Medical Sciences, P20GM130457 (G.A. Hobbs)

Scott and Cynthia McAfee’s Fund for Pancreatic Cancer Precision Medicine (M. Kamgar, T. McFall)

Craig Adelman, the Burkhardt Family Fund, and the Mike Welsh Fund for Pancreatic Cancer Research (M. Kamgar, T. McFall, Y.D. Seo, N.K. Lytle)

Joel and Arlene Lee Chair of Pancreatic Cancer Research (M. Kamgar)

## DATA AVAILABILITY

All data generated during this study are included in this published article [and its supplementary information files]. Any non-commercial reagents generated by the authors are available upon request. Genomic profiling for the patient was performed by Tempus, a CLIA-certified lab. Accessing data beyond what was published in the current study would require a data usage agreement with the corresponding commercial companies. Bioinformatic data generated in this study are publicly available in SRA under BioProject accession number PRJNA1459483.

## CODE AVAILABILITY

Custom Python code used for image preprocessing, ResNet-50–based feature extraction, and cosine similarity-based degradation quantification of toroid images is available as a Python package on GitHub (github.com/tmcfall85/McFallLab/blob/main/mcw_utils/donut_quant/README.md). The package depends on Python (v3.12.7), the Python Imaging Library (v10.4.0), PyTorch (v2.6.0+cu126), the Hugging Face transformers library (v4.48.2) with the pretrained ResNet-50 model, and NumPy (v1.26.4). Bioinformatics analysis was performed by Novogene using their standard pipeline (BWA/GTK/Annovar); variants with exonic or splicing functions were cross-referenced against OncoKB and filtered for at least a likely oncogenic effect. All custom scripts are available at github.com/eke-osumc/McFall-VanillaScoop.

---

**Figure S1.**
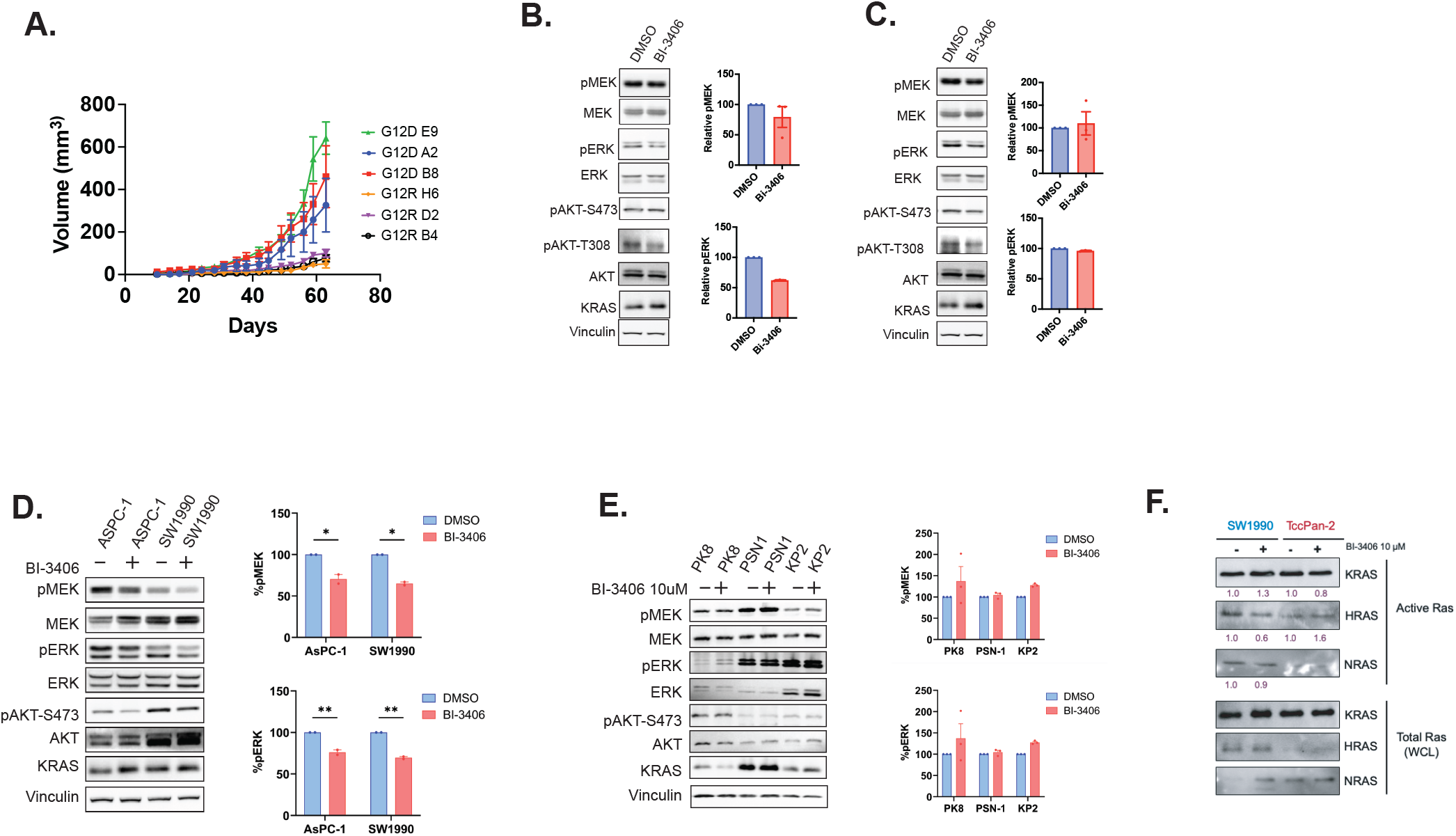
Role of KRASMur .sos axis in MAPK signaling. **A**. Three independent PANC-1 G12D and G12R clones were additionally evaluated for in vivo growth kinetics following xenografting of isogenic cell lines into NSG mice (n = 5 per group), with tumor growth monitored over 58 days. Data points represent mean values, and error bars indicate standard deviation points represent mean values, and error bars indicate standard deviation. **B**. Left: PANC-1 KRASG^120^ cells (in serum free media) were treated with vehicle (DMSO) or 10 µM BI-3406 for 12 hours. Cells were lysed, and proteins were resolved by SDS-PAGE and immunoblotted for RAS pathway proteins and their phosphorylated (activated) forms. Right: Quantification of phosphorylated MEK (pMEK) and phosphorylated ERK (pERK) from three independent experiments shown in panel A. **C**. Left: PANC-1 KRASG^12^R cells (in serum free media) were treated with vehicle (DMSO) or 10 µM BI-3406 for 12 hours. Cells were lysed, and proteins were resolved by SDS-PAGE and immunoblotted for RAS pathway proteins and their phosphorylated (activated) forms. Right: Quantification of pMEK and pERK from three independent experiments shown in panel B. **D**. Left: Library of KRASG^120^ PDAC cells (in complete media) were treated with vehicle (DMSO) or 10 **µM** BI-3406 for 2 hours. Cells were lysed, and proteins were resolved by SDS-PAGE and immunoblotted for RAS pathway proteins and their phosphorylated (activated) forms. Right: Quantification from three separate experiments for pMEK and pERK. **E**. Library of KRASG^12R^ PDAC cells (in complete media) were treated with vehicle (DMSO) or 10 **µM** BI-3406 for 2 hours. Cells were lysed, and proteins were resolved by SDS-PAGE and immunoblotted for RAS pathway proteins and their phosphorylated (activated) forms. Right: Quantification from three separate experiments for pMEK and pERK. **F**. SW1990 (KRASG^120)^ and TccPan-2 (KRASG^12R)^ cells were maintained in 10% FBS, serum-starved overnight, and treated with BI-3406 (10 **µM)** for 2 hours. Active RAS isoforms were isolated by RBD pulldown from whole-cell lysates. Precipitated fractions and inputs were resolved by SDS-PAGE and immunoblotted for KRAS, HRAS, and NRAS.

**Figure S2.**
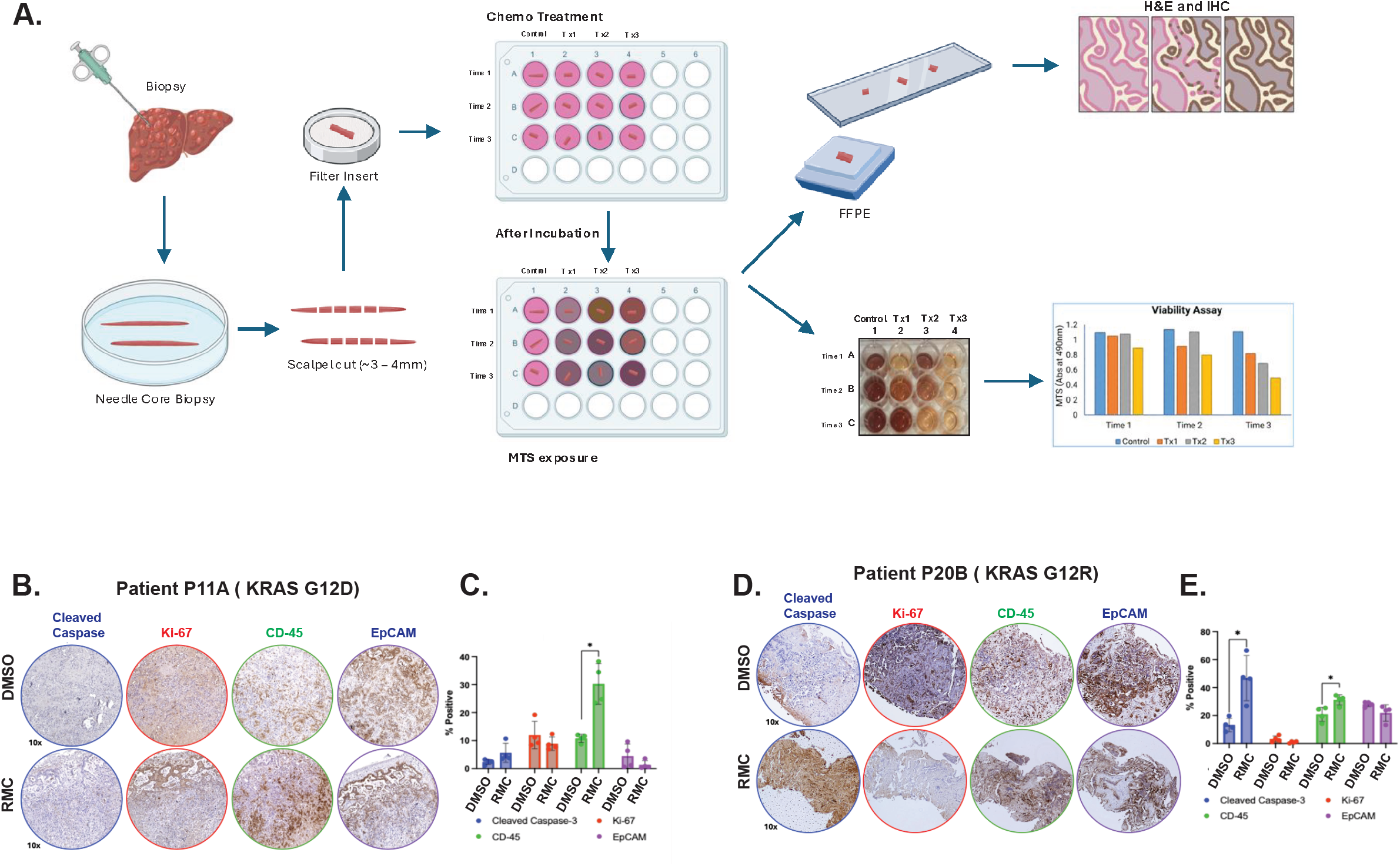
Ex-vivo organotypic slice culture methodology. **A**. Schematic representation of methodology and sample development process for Organotypic Ex-vivo culture **B**. Patient P11A (KRASG120) tumor tissue was cultured ex vivo and treated with RMC-6236 for 48 hours. Fixed samples were immunostained for cleaved caspase-3, Ki-67, CD45, and EpCAM. **C**. Quantification of IHC staining from three independent sections of patient P11A tumor. **D**. Patient P20B (KRASG^12^R) tumor tissue was cultured ex vivo and treated with RMC-6236 for 48 hours. Fixed samples were immunostained for cleaved caspase-3, Ki-67, CD45, and EpCAM. **E**. Quantification of IHC staining from three independent sections of patient P20B tumor. All experiments were performed at least three times for statistical rigor. Statistical differences were calculated by two-tailed unpaired t-test and P values are indicated (for two groups). For experiments with three or more groups statistical comparisons were calculated by one-way ANOVA followed by post hoc Tukey’s test for multiple comparisons, and P values are indicated in the panels. Each experiment is representative of three experiments; quantifications represent mean and error bars represent standard deviations from three separate experiments

**Figure S3.**
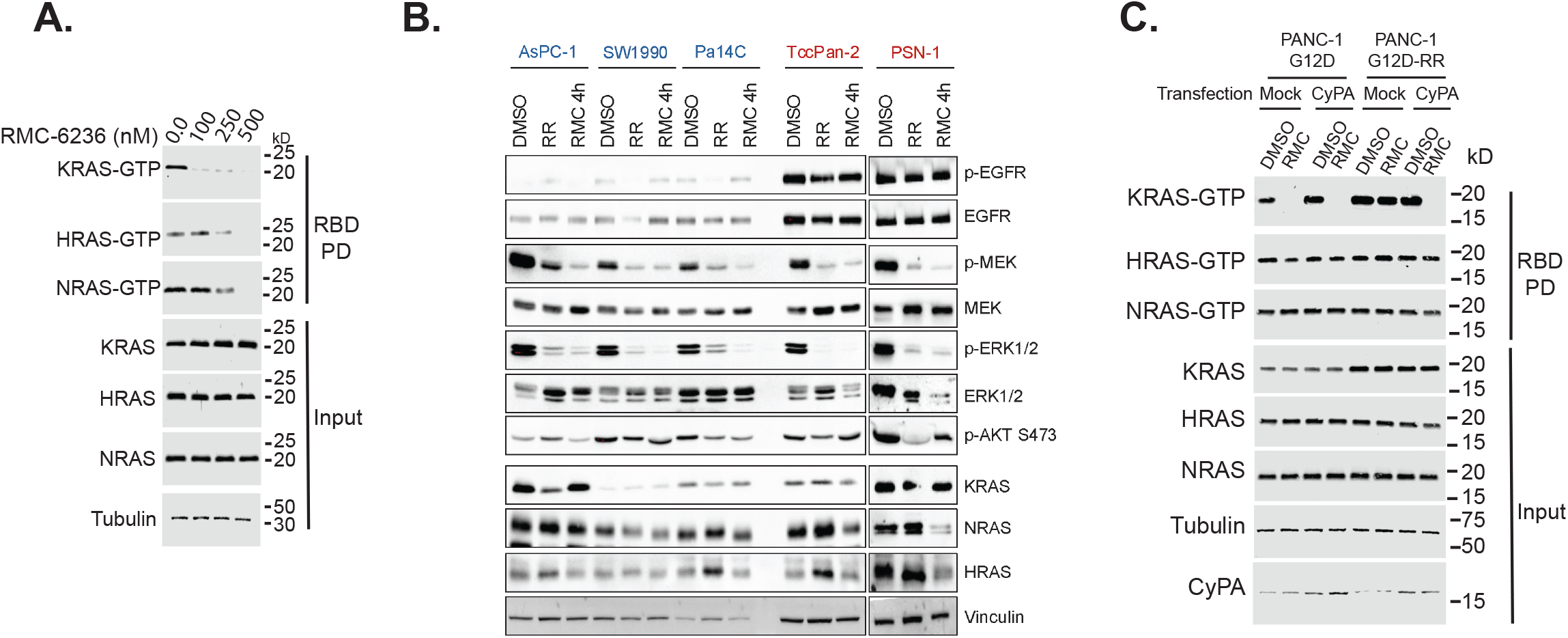
RAS inhibitor resistance. **A**. Active RAS pulldown assay with higher concentrations of daraxonrasib. **B**. ASPC-1, SW1990, Pa14C, TccPan-2, and PSN-1 cells were cultured in media containing RMC-7977 (10 nM) for 6 weeks, in standard media for respective cell lines, or treated with 10 nM RMC-7977 for 4 hours. Lysates were resolved by SDS-PAGE and immunoblotted for proteins in the RAS pathway as indicated. KRASG^1^2D cell lines are noted in blue and G12R in red. **C**. PANC-1 G12D and PANC-1 G12D-RR cells we either transfected with empty vector or CyPA PcDNA, RBD pulldowns were performed after 24 hour treatment with RMC-6236 (80nM) and pulldown product and input lysates were resolved by SDS-PAGE and blotted for respective proteins.

**Fig. S4.**
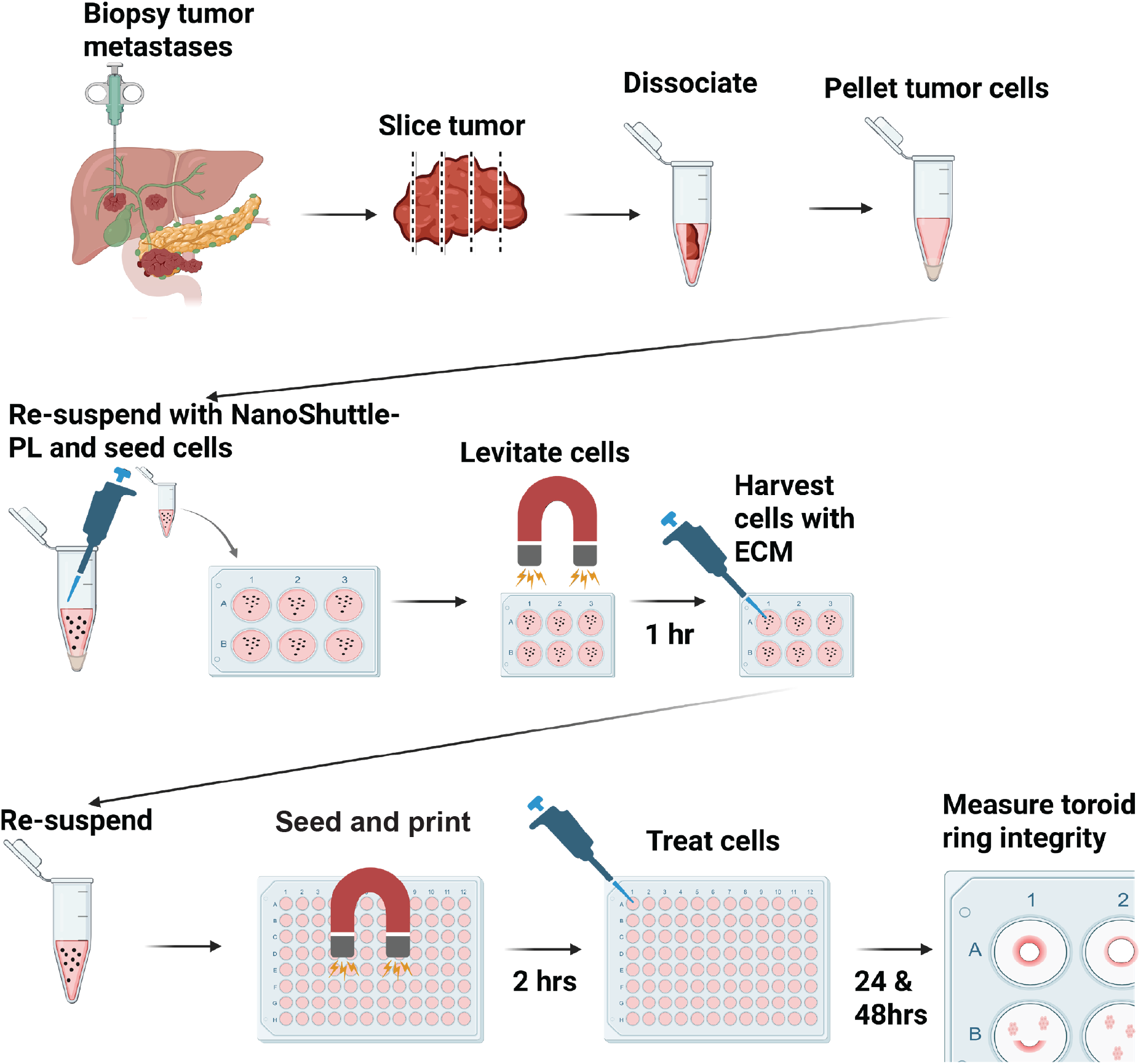
Patient Tumor Bioprinting. Patient-derived pancreatic tumor samples were obtained via biopsy or resection by the LaBahn Pancreas Group at the Medical College of Wisconsin. Tumor tissue was mechanically dissected and enzymatically dissociated in the laboratory. Dissociated cells were pelleted, resuspended in human pancreatic media supplemented with NanoShuttle-PL, and plated in ultra-low attachment 6-well plates. Cells were levitated using a levitation drive for 1 hour to facilitate three-dimensional assembly. The resulting cell-ECM constructs were collected, resuspended, and bioprinted using a ring drive for 2 hours. Following structure formation and quality assessment, organoids were treated with indicated compounds. Images were acquired immediately post-treatment (30 min) and at subsequent timepoints as indicated.

**Figure S5.**
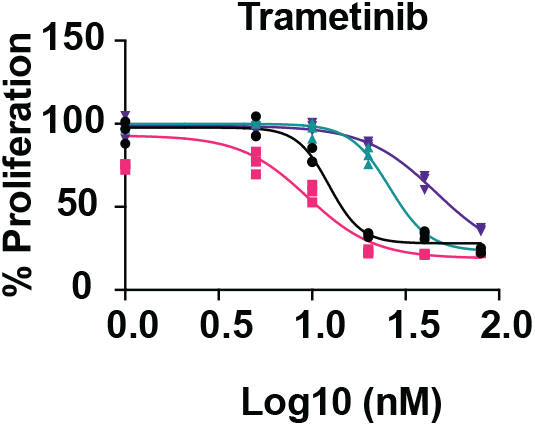
Sensitivity to trametinib. PANC-1 KRAS^G12R^, PANC-1 KRAS^G12D^ and their resistant derivatives (-RR) were treated with increasing concentrations of trametinib for 5 days, and cell proliferation was determined by MTT assay.

**Figure S6.**
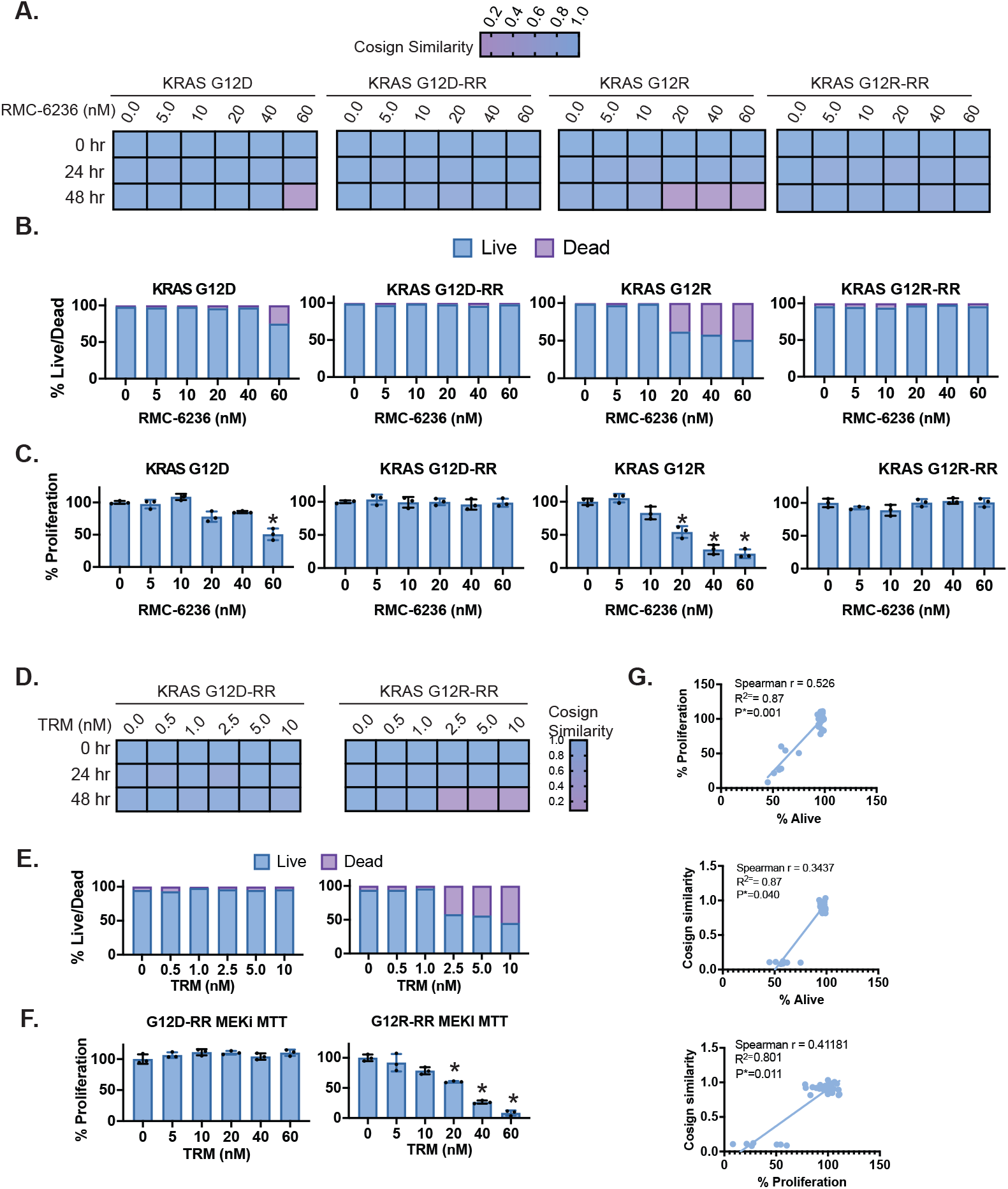
Characterization of Toroid integrity by orthogonal methods. (A)Cosine similarity score for each toroid treatment group in Fig. 7A. (B)After imaging toroids at the 48 h time point, toroids were disrupted and cells were subjected to a trypan blue exclusion assay to determine the percentage of live and dead cells. (C)The remaining dissociated cells were subjected to a CellTiter-Glo assay to determine relative proliferation compared to the DMSO control at the 48 h time point shown in (A). (D)Cosine similarity score for each toroid treatment group in Fig. 7F. (E)After imaging toroids at the 48 h time point (Fig. 7F), toroids were disrupted and cells were subjected to a trypan blue exclusion assay to determine the percentage of live and dead cells. (F)The remaining dissociated cells were subjected to a CellTiter-Glo assay to determine relative proliferation compared to the DMSO control at the 48 h time point shown in (D). (G)Pearson correlation analysis was performed across the metrics from the three assays to assess their concordance.

## REFERENCES

1. Yousef A, Yousef M, Chowdhury S, Abdilleh K, Knafl M, Edelkamp P, et al. Impact of KRAS mutations and co-mutations on clinical outcomes in pancreatic ductal adenocarcinoma. Npj Precis Oncol. 2024;8:27.

2. Jiang J, Jiang L, Maldonato BJ, Wang Y, Holderfield M, Aronchik I, et al. Translational and Therapeutic Evaluation of RAS-GTP Inhibition by RMC-6236 in RAS-Driven Cancers. Cancer Discov. 2024;14:994–1017.

3. Cregg J, Edwards AV, Chang S, Lee BJ, Knox JE, Tomlinson ACA, et al. Discovery of Daraxonrasib (RMC-6236), a Potent and Orally Bioavailable RAS(ON) Multi-selective, Noncovalent Tri-complex Inhibitor for the Treatment of Patients with Multiple RAS-Addicted Cancers. J Med Chem. 2025;68:6064–83.

4. Holderfield M, Lee BJ, Jiang J, Tomlinson A, Seamon KJ, Mira A, et al. Concurrent inhibition of oncogenic and wild-type RAS-GTP for cancer therapy. Nature. 2024.

5. Filis P, Salgkamis D, Matikas A, Zerdes I. Breakthrough in RAS targeting with pan-RAS (ON) inhibitors RMC-7977 and RMC-6236. Drug Discov Today. 2025;30:104250.

6. O’Reilly EM, Wainberg ZA, Hendifar AE, Borad MJ, Pietrantonio F, Pant S, et al. Daraxonrasib or chemotherapy in previously treated metastatic pancreatic cancer. N Engl J Med. 2026.

7. Olivier T, Haslam A, Prasad V. Post-progression treatment in cancer randomized trials: a cross-sectional study of trials leading to FDA approval and published trials between 2018 and 2020. BMC Cancer. 2023;23:448.

8. Honkala A, Malhotra SV, Kummar S, Junttila MR. Harnessing the predictive power of preclinical models for oncology drug development. Nat Rev Drug Discov. 2022;21:99–114.

9. Isermann T, Sers C, Der CJ, Papke B. KRAS inhibitors: resistance drivers and combinatorial strategies. Trends Cancer. 2025;11:91–116.

10. Awad MM, Liu S, Rybkin II, Arbour KC, Dilly J, Zhu VW, et al. Acquired Resistance to KRASG12C Inhibition in Cancer. N Engl J Med. 2021;384:2382–93.

11. Adachi Y, Ito K, Hayashi Y, Kimura R, Tan TZ, Yamaguchi R, et al. Epithelial-to-Mesenchymal Transition is a Cause of Both Intrinsic and Acquired Resistance to KRAS G12C Inhibitor in KRAS G12C-Mutant Non-Small Cell Lung Cancer. Clin Cancer Res. 2020;26:5962–73.

12. Stites EC, Trampont PC, Ma Z, Ravichandran KS. Network analysis of oncogenic Ras activation in cancer. Science. 2007;318:463–7.

13. Kamgar M, Hobbs GA, Davidson R, Dorbin D, Herrera J, Langenheim JF, et al. Mutant and Wild-type RAS Crosstalk and Stoichiometric Deficiencies are Determinants of Sensitivity to Targeted Therapies in KRASG12R Pancreatic Ductal Adenocarcinoma. Cancer Res. 2026.

14. McFall T, Stites EC. Identification of RAS mutant biomarkers for EGFR inhibitor sensitivity using a systems biochemical approach. Cell Rep. 2021;37:110096.

15. McFall T, Schomburg NK, Rossman KL, Stites EC. Discernment between candidate mechanisms for KRAS G13D colorectal cancer sensitivity to EGFR inhibitors. Cell Commun Signal. 2020;18:1–7.

16. McFall T, Diedrich JK, Mengistu M, Littlechild SL, Paskvan KV, Sisk-Hackworth L, et al. A systems mechanism for KRAS mutant allele–specific responses to targeted therapy. Sci Signal. 2019;12.

17. McFall T, Trogdon M, Guizar AC, Langenheim JF, Sisk-Hackworth L, Stites EC. Co-targeting KRAS G12C and EGFR reduces both mutant and wild-type RAS-GTP. NPJ Precis Oncol. 2022;6:86.

18. He K, Zhang X, Ren S, Sun J. Deep residual learning for image recognition. 2016. p.770–8.

19. Hunter JC, Manandhar A, Carrasco MA, Gurbani D, Gondi S, Westover KD. Biochemical and Structural Analysis of Common Cancer-Associated KRAS Mutations. Mol Cancer Res. 2015;13:1325–35.

20. McIntyre CA, Grimont A, Park J, Meng Y, Sisso WJ, Seier K, et al. Distinct clinical outcomes and biological features of specific KRAS mutants in human pancreatic cancer. Cancer Cell. 2024;42:1614–1629.e5.

21. Burge RA, Le Roux O, Popow O, Spadafora VK, Rajesh C, Adair SJ, et al. KRASG12R-Mutant Pancreatic Cancer Features Limited ERK/MAPK Transcriptional Activity and a Distinctive Tumor Microenvironment. Cancer Res. 2026.

22. Hobbs GA, Baker NM, Miermont AM, Thurman RD, Pierobon M, Tran TH, et al. Atypical KRASG12R mutant is impaired in PI3K signaling and macropinocytosis in pancreatic cancer. Cancer Discov. 2020;10:104–23.

23. Cuevas-Navarro A, Pourfarjam Y, Hu F, Rodriguez DJ, Vides A, Sang B, et al. Pharmacological restoration of GTP hydrolysis by mutant RAS. Nature. 2025;637:224–9.

24. Yang MH, Tran TH, Hunt B, Agnor R, Johnson CW, Shui B, et al. Allosteric Regulation of Switch-II Domain Controls KRAS Oncogenicity. Cancer Res. 2023;83:3176–83.

25. Stites EC, Trampont PC, Ma Z, Ravichandran KS. Network analysis of oncogenic Ras activation in cancer. Science. 2007;318:463–7.

26. Hood FE, Sahraoui YM, Jenkins RE, Prior IA. Ras protein abundance correlates with Ras isoform mutation patterns in cancer. Oncogene. 2023;42:1224–32.

27. Mageean CJ, Griffiths JR, Smith DL, Clague MJ, Prior IA. Absolute Quantification of Endogenous Ras Isoform Abundance. PloS One. 2015;10:e0142674.

28. Prior IA, Hood FE, Hartley JL. The Frequency of Ras Mutations in Cancer. Cancer Res. 2020;80:2969–74.

29. Conroy T, Desseigne F, Ychou M, Bouché O, Guimbaud R, Bécouarn Y, et al. FOLFIRINOX versus gemcitabine for metastatic pancreatic cancer. N Engl J Med. 2011;364:1817–25.

30. Souza GR, Molina JR, Raphael RM, Ozawa MG, Stark DJ, Levin CS, et al. Three-dimensional tissue culture based on magnetic cell levitation. Nat Nanotechnol. 2010;5:291–6.

31. Jaganathan H, Gage J, Leonard F, Srinivasan S, Souza GR, Dave B, et al. Three-dimensional in vitro co-culture model of breast tumor using magnetic levitation. Sci Rep. 2014;4:6468.

32. Hou S, Tiriac H, Sridharan BP, Scampavia L, Madoux F, Seldin J, et al. Advanced development of primary pancreatic organoid tumor models for high-throughput phenotypic drug screening. SLAS Discov. 2018;23:574–84.

33. Haisler WL, Timm DM, Gage JA, Tseng H, Killian T, Souza GR. Three-dimensional cell culturing by magnetic levitation. Nat Protoc. 2013;8:1940–9.

34. Baillargeon P, Shumate J, Hou S, Fernandez-Vega V, Marques N, Souza G, et al. Automating a magnetic 3D spheroid model technology for high-throughput screening. SLAS Technol. 2019;24:420–8.

35. Yoshida T, Kakegawa J, Yamaguchi T, Hantani Y, Okajima N, Sakai T, et al. Identification and characterization of a novel chemotype MEK inhibitor able to alter the phosphorylation state of MEK1/2. Oncotarget. 2012;3:1533.

36. Tran KA, Cheng MY, Mitra A, Ogawa H, Shi VY, Olney LP, et al. MEK inhibitors and their potential in the treatment of advanced melanoma: the advantages of combination therapy. Drug Des Devel Ther. 2016;10:43–52.

37. Gruber A, Czejka M, Buchner P, Kitzmueller M, Kirchbaumer Baroian N, Dittrich C, et al. Monitoring of erlotinib in pancreatic cancer patients during long-time administration and comparison to a physiologically based pharmacokinetic model. Cancer Chemother Pharmacol. 2018;81:763–71.

38. Bapiro TE, Richards FM, Goldgraben MA, Olive KP, Madhu B, Frese KK, et al. A novel method for quantification of gemcitabine and its metabolites 2′,2′-difluorodeoxyuridine and gemcitabine triphosphate in tumour tissue by LC–MS/MS: comparison with 19F NMR spectroscopy. Cancer Chemother Pharmacol. 2011;68:1243–53.

39. Moore MJ, Goldstein D, Hamm J, Figer A, Hecht JR, Gallinger S, et al. Erlotinib plus gemcitabine compared with gemcitabine alone in patients with advanced pancreatic cancer: a phase III trial of the National Cancer Institute of Canada Clinical Trials Group. J Clin Oncol. 2007;25:1960–6.

40. Ardalan B, Azqueta JI, England J, Eatz TA. Potential benefit of treatment with MEK inhibitors and chemotherapy in BRAF-mutated KRAS wild-type pancreatic ductal adenocarcinoma patients: a case report. Mol Case Stud. 2021;7:a006108.

41. Kinsey CG, Camolotto SA, Boespflug AM, Guillen KP, Foth M, Truong A, et al. Protective autophagy elicited by RAF→MEK→ERK inhibition suggests a treatment strategy for RAS-driven cancers. Nat Med. 2019;25:620–7.

42. Löhr M, Klöppel G, Maisonneuve P, Lowenfels AB, Lüttges J. Frequency of K-ras mutations in pancreatic intraductal neoplasias associated with pancreatic ductal adenocarcinoma and chronic pancreatitis: a meta-analysis. Neoplasia. 2005;7:17–23.

43. Labadie BW, Obradovic A, Raufi AG, Wasko UN, Tomassoni L, Ge L, et al. Targeting of MEK and Autophagy in Pancreatic Adenocarcinoma and Analysis of Treatment Sensitivity in Preclinical and Clinical Models: MEKiAUTO. JCO Precis Oncol. 2026;10:e2600138.

44. Long S-A, Todd H, Goodhart G, Chang W-H, Amparo AM, Bridgens R, et al. CRISPR-Cas9 Screening Identifies Resistance Mechanisms to KRAS Inhibition in Pancreatic Cancer. Cancer Res. 2026;86:1035–53.

45. Feng J, Hu Z, Xia X, Liu X, Lian Z, Wang H, et al. Feedback activation of EGFR/wild-type RAS signaling axis limits KRASG12D inhibitor efficacy in KRASG12D-mutated colorectal cancer. Oncogene. 2023;42:1620–33.

46. Liu Z, Lenz H-J, Yu J, Zhang L. Differential Response and Resistance to KRAS-Targeted Therapy. Mol Carcinog. 2025;64:1135–48.

47. Ash LJ, Busia-Bourdain O, Okpattah D, Kamel A, Liberchuk A, Wolfe AL. KRAS: Biology, Inhibition, and Mechanisms of Inhibitor Resistance. Curr Oncol. 2024;31:2024–46.

48. Zeiser R, Andrlová H, Meiss F. Trametinib (GSK1120212). In: Martens UM, editor. Small Mol Oncol. Cham: Springer International Publishing; 2018. p.91–100.

49. Hobbs GA, Baker NM, Miermont AM, Thurman RD, Pierobon M, Tran TH, et al. Atypical KRASG12R Mutant Is Impaired in PI3K Signaling and Macropinocytosis in Pancreatic Cancer. Cancer Discov. 2020;10:104–23.

50. Knox JJ, O’Kane G, King D, Laheru D, Habowski AN, Yu K, et al. PASS-01: Randomized Phase II Trial of Modified FOLFIRINOX Versus Gemcitabine/Nab-Paclitaxel and Molecular Correlatives for Previously Untreated Metastatic Pancreatic Cancer. J Clin Oncol. 2025;JCO-25-00436.

51. Gueiderikh A, Tarabay A, Abdelouahab M, Smolenschi C, Tanguy M, Valery M, et al. Pancreatic adenocarcinoma third line systemic treatments: a retrospective cohort study. BMC Cancer. 2024;24:272.

52. Fan M, Deng G, Ma Y, Si H, Wang Z, Dai G. Survival outcome of different treatment sequences in patients with locally advanced and metastatic pancreatic cancer. BMC Cancer. 2024;24:67.

53. Kim B, Ahn J, Jung JH, Jung K, Lee J-C, Hwang J-H, et al. Efficacy of the third-line chemotherapy in patients with advanced pancreatic cancer. J Clin Oncol. 2023;41:711–711.

